# ADAP’s intrinsically disordered region is an actin sponge regulating T cell motility

**DOI:** 10.1101/2021.12.14.472590

**Authors:** Nirdosh Dadwal, Janine Degen, Jana Sticht, Tarek Hilal, Tatjana Wegner, Peter Reichardt, Ruth Lyck, Michael Abadier, Miroslav Hons, Charlie Mix, Benno Kuropka, Heike Stephanowitz, Fan Liu, Burkhart Schraven, Christoph Wülfing, Stefanie Kliche, Christian Freund

## Abstract

Intrinsically disordered proteins (IDPs) play a vital role in biological processes that rely on transient molecular compartmentation^1^. In T cells, the dynamic switching between migration and adhesion mandates a high degree of plasticity in the interplay of adhesion and signaling molecules with the actin cytoskeleton^2,3^. Here, we show that the N-terminal intrinsically disordered region (IDR) of adhesion- and degranulation-promoting adapter protein (ADAP) acts as a multipronged scaffold for G- and F-actin, thereby promoting actin polymerization and bundling. Positively charged motifs, along a sequence of at least 200 amino acids, interact with both longitudinal sides of G-actin in a promiscuous manner. These polymorphic interactions with ADAP become constrained to one side once F-actin is formed. Actin polymerization by ADAP acts in synergy with a capping protein but competes with cofilin. In T cells, ablation of ADAP impairs adhesion and migration with a time-dependent reduction of the F-actin content in response to chemokine or T cell receptor (TCR) engagement. Our data suggest that IDR-assisted molecular crowding of actin above the critical concentration defines a new mechanism to regulate cytoskeletal dynamics. The principle of IDRs serving as molecular sponges to facilitate regulated self-assembly of filament-forming proteins might be a general phenomenon.

---

T cells are migratory cells that need to switch to adhesion in the extravasation from blood vessels and subsequent interaction with antigen presenting cells. Chemokine receptors trigger the directed migration of T cells into secondary lymphatic organs (SLOs) while the engagement of the TCR via peptide-loaded MHCs on antigenpresenting cells (APC) leads to the formation of a specialized cell – cell junction^4–6^. Changes in integrin affinity and avidity are at the core of switching cellular behavior and are regulated by the scaffolding protein ADAP^7,8^. ADAP does not interact directly with the TCR, chemokine receptors or integrins, yet it constitutes the common signaling component connecting these receptors. Two atypical hSH3 domains confer weak lipid binding^9^ and activation-dependent phosphorylation of the interdomain linker engages SH2 domain-containing kinases and adaptor proteins^10–15^. While ADAP was found to co-localize with actin-rich patches in lamellipodia of Jurkat cells^16^ and is linked to the actin regulators Ena/VASP and Nck^17,18^, it’s conceivable role as a driver of actin dynamics has not been addressed.

## Actin-dependent cellular processes are compromised in ADAP^-/-^ cells

We hypothesized that actin regulation by ADAP might explain critical features of T cell function upon chemokine or TCR stimulation. We have previously shown that ADAP is crucial for T cell homing *in vivo*^19^. In 2-Photon microscopy experiments with adoptively transferred and differentially labeled WT and ADAP^-/-^ T cells, the interaction of ADAP^-/-^ T cells with the intranodal vessel wall of the inguinal lymph node *in vivo* was attenuated (Fig. 1a). The number of stably arrested T cells was reduced, while the number of transient encounters was enhanced (Fig. 1b). In shear flow studies on Primary Mouse Brain Microvascular Endothelial Cells (pMBMECs) *ex vivo* loss of ADAP impaired stable T cell arrest (Extended Data Fig. 1a,b, Supplementary Movie 1). ADAP-deficiency led to a reduced frequency of crawling T cells and an enhanced detachment rate (Fig. 1c). Moreover, ADAP^-/-^ T cells were less polarized as reflected by a reduced T cell length-to-width ratio (Extended Data Fig. 1c). This polarization defect correlated with reduced F-actin polymerization at multiple time points after chemokine stimulation (Fig. 1d and Extended Data Fig. 1d), indicating that compromised F-actin dynamics is involved in the reduced homing capacity of ADAP^-/-^ T cells. Similarly, the increase in F-actin content upon triggering of the TCR was attenuated in ADAP^-/-^ T cells (Fig. 1e). The T cell area upon spreading on a cover slip coated with a stimulating anti-TCR antibody was greatly diminished (Extended Data Fig. 1e). Consistent with a role of ADAP in F-actin turnover, ADAP localized to the periphery of the cellular interface during the first minutes of cognate activation of primary murine T cells by antigen presenting cells (Extended Data Fig. 1f). This distribution was similar to that of F-actin, the core actin regulatory machinery exemplified by CapZ and cofilin^20^ and of ADAP’s constitutive binding partner SKAP55 (Extended Data Fig. 1f). In Jurkat T cells, ADAP colocalized with F-actin within the lamellipodia with a small but significant proportion of ADAP found at the rim, where Factin nucleation is thought to be initialized (Fig. 1f). In summary, the F-actin-dependent processes of T cell polarization, migration, crawling and adhesion are compromised in the absence of ADAP.

**Fig. 1:**
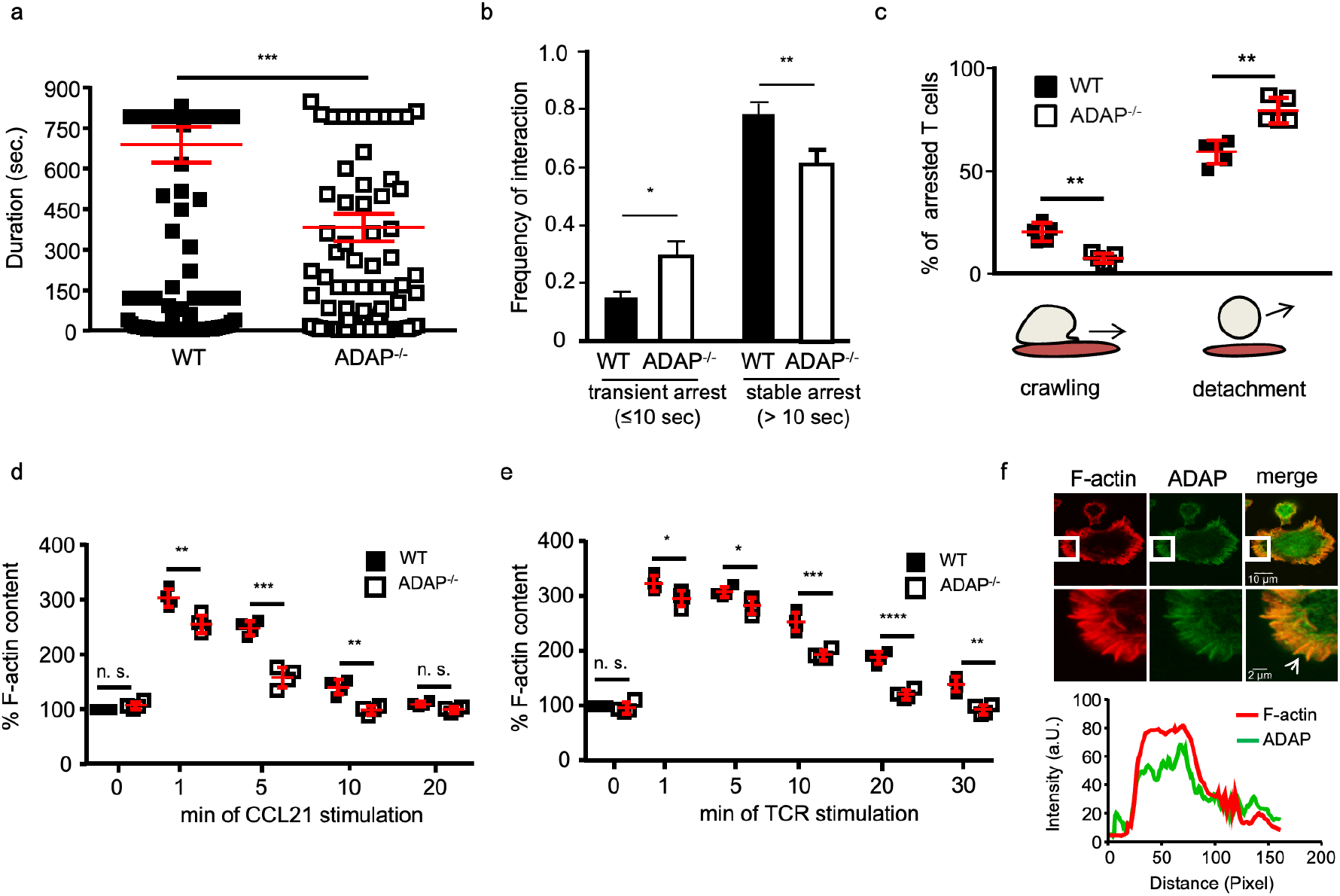
ADAP compromises F-actin polymerization and cytoskeleton-dependent functions in T cells. (a) 2-Photon microscopic analysis of the duration of the interaction of CFSE-labeled WT and DDAO-SE-labeled ADAP^-/-^ T cells with the intranodal vessel wall *in vivo* ((n=3 experiments, n=111 WT T cells, n = 126 ADAP^-/-^ T cells,**p<0.01 (b) Analysis of frequency of interaction with the intranodal vessel within 10 sec (transient arrest) or longer than 10 sec (stable arrest) (n=3 experiments, n=111 WT T cells, n = 126 ADAP^-/-^ T cells, *p<0.01,**p<0.01 ***p<0.001). (c) CMTMR-labeled WT and blueCMAC-labeled ADAP^-/-^ T cells were analyzed on CCL21-coated TNF-α-stimulated pMBMECs under shear flow (1.5 dyn/cm^2^) for 30 min. The percentages of T cell crawling and detachment were calculated from the pool of initially arrested WT and ADAP^-/-^ T cells (see Extended Figure 1b) (n=2, 5 films, ***p<0.001) (d-e). Isolated naïve WT and ADAP^-/-^ T cells were either stimulated with (d) CCL21 or (e) anti-CD3 antibodies for the indicated times. T cells were permeabilized, fixed, and stained with FITC-phalloidin. The F-actin content was measured by flow cytometry. The mean fluorescence intensity of untreated WT T cells was set to 100%. (n=4, *p<0.05, **p<0.01, ***p<0.001, ****p<0.0001). (f) Jurkat T cells were settled on slides precoated with anti-CD3 mAbs. The cells were fixed, permeabilized and stained with anti-ADAP mAbs in combination with anti-rat IgG-FITC (green). F-actin was visualized with TRITC-Phalloidin (red), cells were imaged by confocal microscopy. The squares indicate the areas that were imaged with higher magnifications. The arrow indicates the thin rim of the lamellipodia. (n=2, 25 cells). A profile intensity plot for ADAP (green) and F-actin (red) is depicted.

## The N-terminus of ADAP directly induces actin polymerization

Recombinantly expressed full-length ADAP or a stable ADAP-SKAP55 complex, ADAP’s predominant form in T cells^21^, induced rapid actin polymerization in a dose dependent manner in an *in vitro* F-actin polymerization assay (Fig 2a, Extended Data Fig. 2a). SKAP55 alone, covalently linked to ADAP_340-450_ to stabilize the protein, did not display any activity (Extended Data Fig. 2b). An intrinsically disordered N-terminal fragment of ADAP comprising the first 381 amino acids (ADAP_1-381_) showed comparable levels of actin polymerization to full-length ADAP (Fig. 2b) while ADAP fragments ADAP_1-245_ and ADAP_1-200_ maintained only partial activity (Extended data Fig. 2c, d). No polymerization was observed for an ADAP C-terminal fragment (ADAP_486-783_) or its SKAP binding region (ADAP_340-486_) (Extended data Fig. 2c,d). Similarly, protein fragments ADAP1-100, ADAP_100-200_ and ADAP_240-340_ were unable to induce actin polymerization (Extended data Fig. 2c,d).

**Fig. 2:**
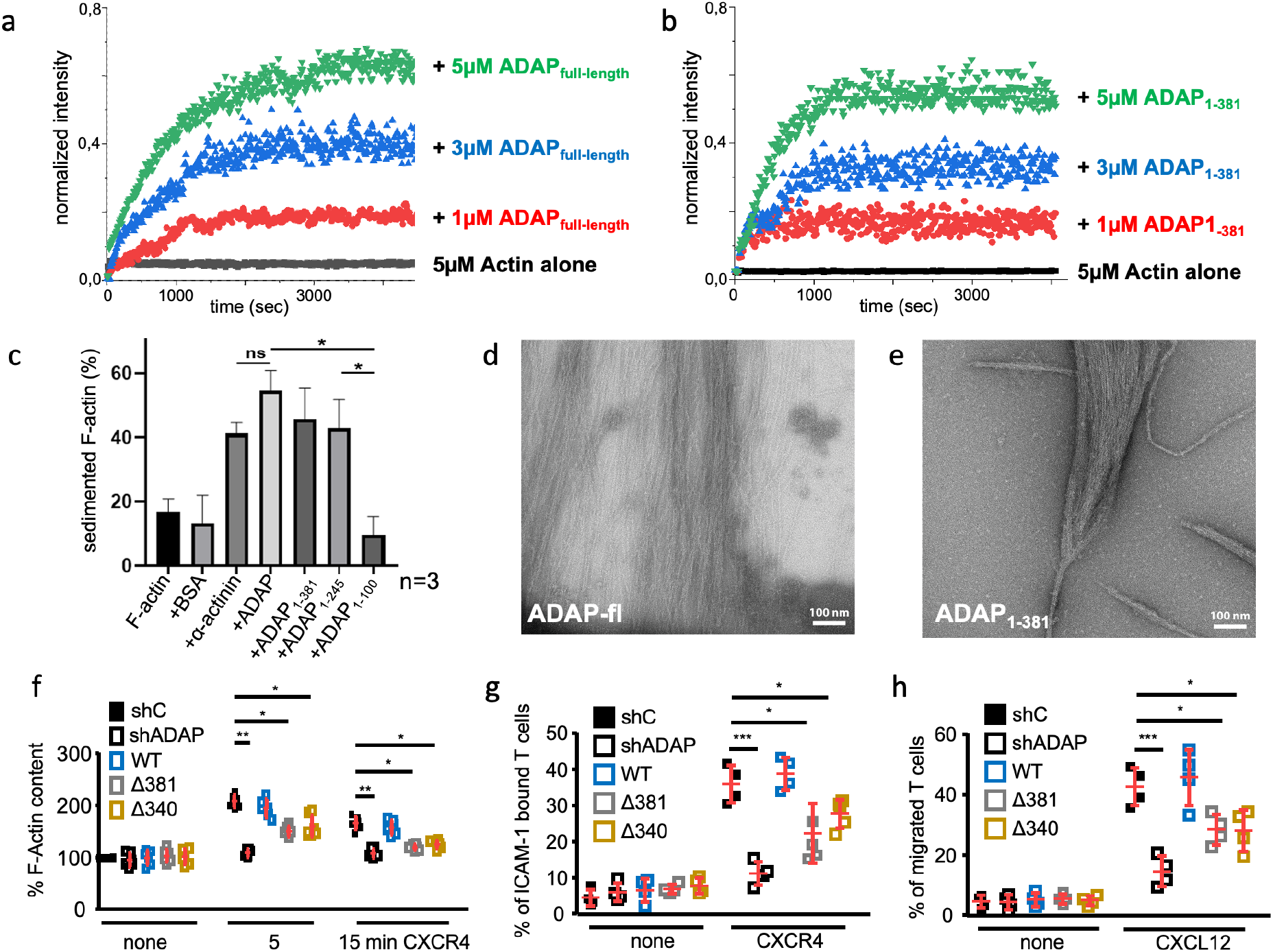
ADAP polymerizes and bundles actin *in vitro.* (a,b) Polymerization curves representing the change in fluorescence intensity of pyrene actin (measured at an interval of 10 seconds, as a result of adding different concentrations (1μM, 3μM and 5μM) of (a) ADAP full-length and (b) ADAP1-381 to 5μM of pyrene-actin. (c) Results of F-actin bundling assay for different ADAP constructs according to the analyzed SDS-gels shown in Extended Data Fig. 4a. (d,e) Negative stain EM pictures of (d) actin polymerized in the presence of ADAP-fl or e) filaments obtained upon incubation of actin with ADAP1-381. (f-h) Functional experiments using Jurkat T cells transfected with suppression/re-expression constructs of ADAP for the F-actin content in response to CXCR4 stimulation (f), percentages of bound T cells to ICAM-1 coated wells upon CXCR4 triggering (g) and (h) percentages of migrated T cells in a transwell assay upon CXCL12 treatment. (n=4-6, student t-test *p<0.05,**p<0.01, ***p<0.001).

To determine whether individual motifs play a dominant role in actin polymerization, we generated a series of mutant proteins of ADAP1-381, each containing deletions of 12-31 amino acids harboring lysine-containing motifs enriched in the sequence (Extended Data Fig. 3a). Each of these deletion constructs behaved similar to wildtype protein (Extended Data Fig. 3b). This indicates that a composite effect of several motifs in the N-terminal region confers actin polymerization-promoting properties to ADAP.

## F-actin bundling is promoted by ADAP

The co-localization of ADAP with F-actin within the lamellipodia (Fig. 1f) led us to investigate F-actin bundling properties of ADAP using low speed *in vitro* cosedimentation. F-actin was co-precipitated by ADAP full-length or ADAP1-381, indicating formation of bundles (Fig. 2c and Extended Data Fig. 4a). ADAP1-245 maintained actin bundling activity, while ADAP1-100 did not (Fig. 2c and Extended Data Fig. 4a), suggesting that those regions in ADAP that allow polymerization also enable bundling of actin. In electron micrographs, individual F-actin filaments (Extended Data Fig. 4b) were observed to assemble into dense and irregular sheets of a length of several hundred nanometers in the presence of full-length ADAP and of ADAP1-381 (Fig. 2d,e, Extended Data Fig. 4c,d). The diameter of these bundles ranged from ten to several hundred nm (Extended Data Fig. 4f). In contrast, the ADAP-SKAP55 complex assembled F-actin into highly regular bundles with diameters between 50 and 73 nm (Extended Data Fig. 4e,f), indicating that the intrinsic polymerizing and actin crosslinking activity of the N-terminus of ADAP is becoming more tightly controlled in the context of the ADAP-SKAP55 complex.

To probe functional consequences of depletion of the N-terminal region of ADAP, we combined the knockdown of endogenous ADAP in Jurkat T cells with reconstitution of ADAP_WT_ or ADAP N-terminal deletion constructs either lacking (ADAP_Δ1-381_) or retaining the SKAP binding site (ADAP_Δ1-340_) (Extended Data Fig. 5a-d). Compared to reconstitution with WT protein, the F-actin content was significantly reduced for both N-terminal deletion constructs. Adhesion and migration of these cells in response to chemokine stimulation was impaired (Fig. 2f-h). We conclude that the interaction of ADAP with SKAP55 is not strictly required for chemokine-triggered, actin-dependent elements of T cell function. TCR-induced changes in F-actin content, adhesion and interaction with APCs were similarly defective (Extended Data Fig. 5e-g). Changes in the surface expression of TCR, CXCR4 and LFA-1 did not account for the observed defects (Extended Data Fig 5h). Confirming the well-known role of the ADAP C-terminus as an inside-out signaling node regulating adhesion^8^, the compromised function in response to chemokine and TCR triggering (Fig 2f-h and Extended Data Fig. 5e-g) were more pronounced for cells depleted of full-length ADAP.

## Positively charged motifs in ADAP_1-381_ mediate multivalent actin interactions

In order to identify molecular motifs mediating the interaction between ADAP and actin, we employed NMR spectroscopy as a method to derive atomistic information on interacting epitopes. The ^1^H-^15^N correlation spectrum of ADAP_1-381_ displayed low dispersion, indicative of an intrinsically disordered protein in line with theoretical predictions (Extended Data Fig. 6a, b). We therefore focused on smaller fragments of the N-terminus (ADAP_1-100_ and ADAP_100-200_) that allowed us to obtain resonance assignments and cover the fragment ADAP_1-200_ that shows polymerization activity (Extended Data Fig. 2d). We utilized non-polymerizable G-actin (NP-actin), to selectively identify binary ADAP-actin interactions.

Individual motifs among the assigned residues such as F21/R22, A34/R35, L38/F39/N40 in ADAP_1-100_, L110/K111 and L148/K149 within an LKP and W131 within a PWPP motif in ADAP_100-200_ showed line-broadening and/or significant chemical shift changes (Fig. 3a, b; Extended Data Fig. 6c-f). Lysine cross-linking mass spectrometry using ADAP_1-381_ showed that K36, K106, K111 and K116 in the vicinity of the residues affected in NMR can cross-link to residues K63, K193 and K330 in NP-actin (Fig. 3c and Extended Data Table 1). These lysines are located on three different subdomains of G-actin (Extended Data Fig. 7a). The linear distance between these residues is between 33 and 48 Å. Thus, cross-linking of individual ADAP lysine residues to all three actin lysines can only occur when the interaction is polymorphic or even fuzzy^22^. To reveal possible positioning of ADAP peptides that cover regions affected by NMR on the surface of G-actin, the observed cross-links were used as soft restraints to dock ADAP residues 17-46 and 97-126 to G-actin. Both peptides primarily cover a similar region on the actin surface in a polymorphic manner. While in the models with ADAP peptide 17-46 residue K36 mainly docks to two sites, the positioning of the three lysine residues in the ADAP peptide 97-126 is more random but retains a preference for either of the two longitudinal faces of actin. (Extended Data Fig. 7b). Utilizing ADAP fragments ADAP_1-100_ and ADAP_100-200_ we observed a similar pattern of lysine crosslinks, with K115 in actin as an additional targeted residue (Extended Data Fig. 7c,d). In addition to the motifs identified by NMR, cross-linking-MS experiments revealed that three lysines each within ADAP residues 167-177, 223-239 and 298-319 cross-link to actin residues K63, K115, K193, K330, implying that these regions additionally contribute to the interaction by contacting the same principal interfaces on the actin monomer (Fig. 3c and Extended Data Fig. 7a).

**Fig. 3:**
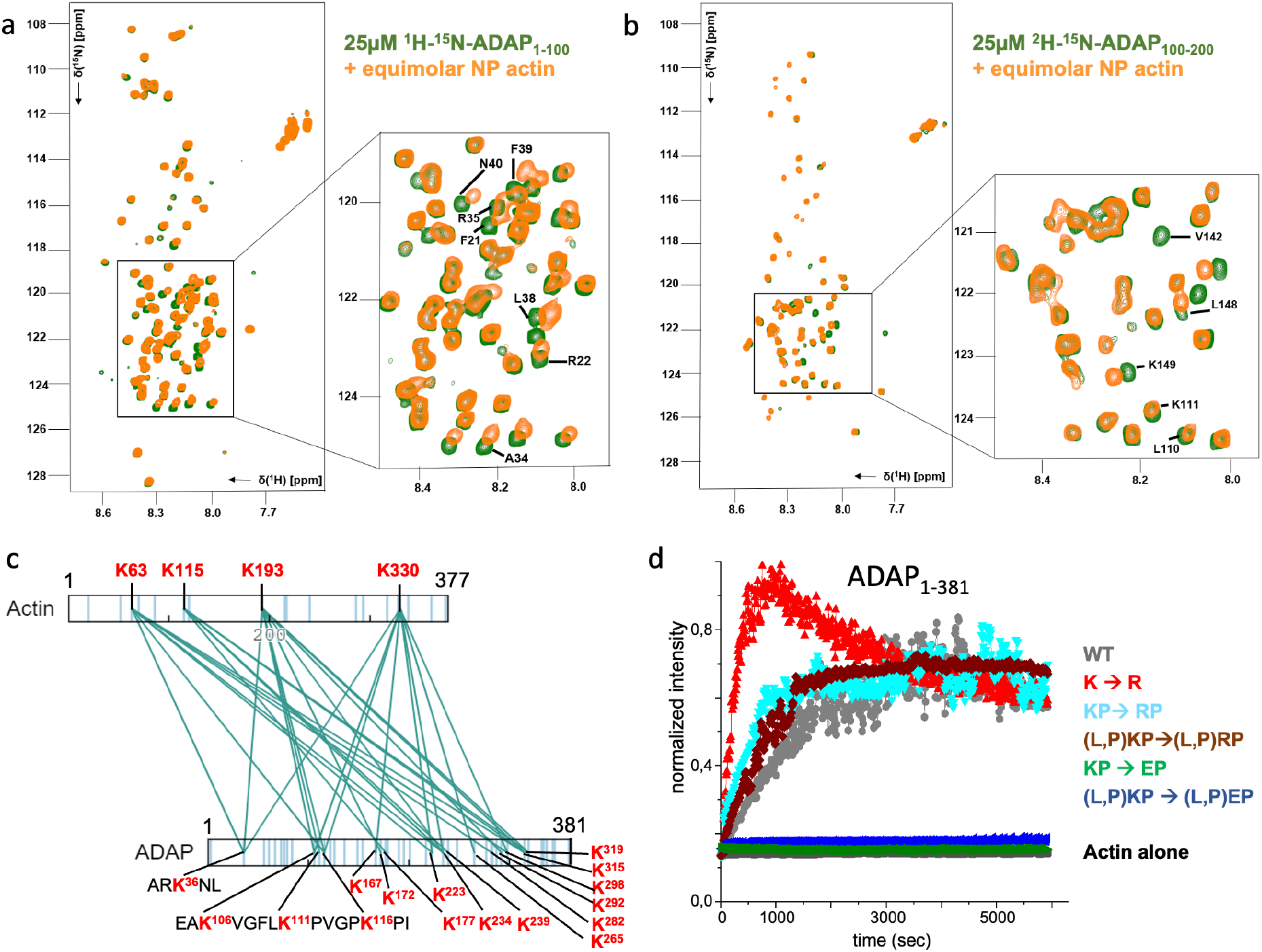
Multiple motifs in ADAP interact promiscuously with actin. ^1^H-^15^N HSQC spectra of ^1^H-^15^N-ADAP_1-100_ (a) and ^2^H-^15^N-ADAP_100-200_ (b) mixed with equimolar amounts of non-polymerizable G-actin. Residues that show either significant chemical shift changes or line broadening are indicated by amino acid type and number c) Lysine residues that were cross-linked between ADAP_1-381_ and NP-actin when incubating stoichiometric amounts of the two proteins. Lysine residues are marked in red and indicated by numbers. (d) Polymerization curves representing the change in fluorescence intensity of pyrene actin (measured at an interval of 10 seconds), as a result of adding 5μM of ADAP_1-381_ or the respective K→R, KP→RP, KP→EP or LKP→LEP/LRP, PKP→PEP/PRP mutants to 5μM of pyrene-actin.

The lysine residues in ADAP_1-381_ are over-represented in KP, LKP or PKP motifs, some of which were affected by NP actin in NMR experiments (e.g. L110, K111, L148, K149). As the individual deletion of stretches containing lysine-rich motifs did not abrogate ADAP’s function in actin polymerization (Extended Data Fig. 3), we probed the importance of charged amino acids in these motifs with ADAP_1-381_ variants where (i) the 17 KP motifs are mutated to either RP or EP and (ii) the 11 LKP and PKP motifs are replaced by LRP/LEP or PRP/PEP. Mutation of K→E in these motifs prevented actin polymerization, supporting the suggestion that the short lysine-containing motifs observed by NMR and cross-linking-MS define sites of interaction (Fig. 3d). In contrast, a K→R substitution in these motifs enhanced actin polymerization. This effect was even more pronounced when all lysines in ADAP_1-381_ are replaced by arginine (Fig. 3d). These data suggest that not only lysine-but also arginine-containing motifs can mediate F-actin polymerization by ADAP. In support, the short motifs F21/R22 and A34/R35 are significantly shifted in the NMR experiments (Fig. 3a, Extended Data Fig. 6c,d). ADAP_1-381_ contains 46 lysines and 11 arginines, suggesting that an optimized polymerization rate of actin, as it is observed by substituting all lysines for arginine, might not be beneficial or regulatable in the context of the cell. In conclusion, F-actin regulation by ADAP is mediated by multivalent, transient and polymorphic interactions of numerous individual sites in ADAP. We suggest that such a situation enhances the local concentration of G-actin in the vicinity of ADAP to a concentration that allows effective F-actin filament formation and elongation.

## F-actin binding by ADAP_1-381_ cooperates with CapZ and competes with cofilin

To further define ADAP’s role in interacting with F-actin, we determined spectral changes in NMR indicative of complex formation. When adding G-actin to ADAP_1-381_, rapid precipitation was seen in the NMR tube (Extended Data Fig. 8a) with an accompanying loss of signal for most ADAP residues. This indicates that the formed actin filaments are bound to the N-terminal part of ADAP, leading to a slowly tumbling polymer containing both proteins. Since line-broadening prevented further structural analysis by NMR, we used cross-linking mass spectrometry (XL-MS) with pre-formed F-actin. Of the three to four sites in G-actin identified to cross-link with ADAP-fragments (Fig. 3c and Extended Data Fig. 7d,e) only two were observed in these experiments, K63 and K330 (Fig. 4a). This appears reasonable as actin residues K115 and K193 are buried in the actin filament (Extended Data Fig. 8b). The cross-linking lysines K63 and K330 are on opposite ends of the actin monomer with a linear distance of 47Å in between them, but they become spatial neighbors in the filament (Extended Data Fig. 8b). Since for several other sites in ADAP_1-381_ (Fig. 4a) simultaneous cross-links to K63 and K330 are detected, this actin dimer interface seems a preferred site of interaction.

**Fig. 4.**
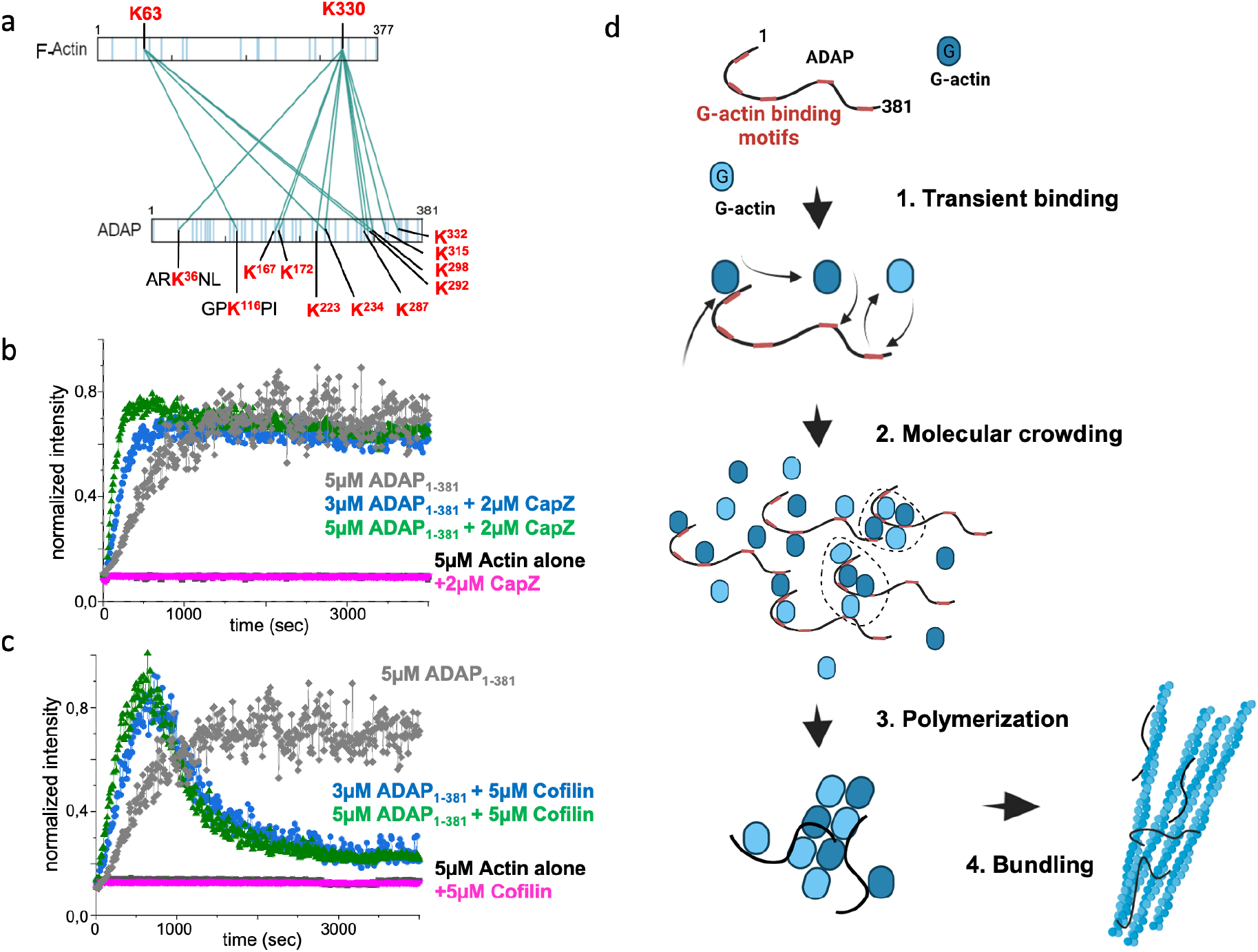
F-actin binding of ADAP. a) Inter-molecular cross-linked lysine residues when incubating stoichiometric amounts of ADAP_1-381_ and pre-polymerized actin shows lysines in ADAP to cross-link to actin K63 and K330. b) Polymerization activity of different concentrations of ADAP_1-381_ (3μM (blue) and 5μM (grey and green)) in the absence (grey) or presence of 2μM CapZ (blue and green) added to 5μM of pyrene-actin c) Polymerization activity of different concentrations of ADAP_1-381_ (3μM (blue) and 5μM (grey and green)) in the absence (grey) or presence of 5μM cofilin added to 5μM of pyrene-actin (blue and green). d) Model for the mode of action of ADAP: 1. The interaction between G-actin and the motifs on ADAP is transient, which means that actin molecules continuously associate and dissociate from the motifs. 2. When multiple molecules of ADAP come together, the continuous binding and un-binding of actin to these motifs on ADAP increases the local concentration of actin. This eventually leads to the formation of a stable actin nucleus (indicated under the dotted circle) initiating polymerization. 3. On the filament ADAP remains bound and polymerizes actin from the pointed end. 4. Once filaments are formed, ADAP will not only bind along the individual actin filaments but also lead to intermolecular binding and thus bundling.

Therefore, we used K63 and K330 from different actin monomers as soft restraints to dock ADAP peptides 17-46 and 97-126 to actin dimers. Individual motifs consistently docked into the groove formed at the interface between two actin monomers (Extended Data Fig. 8c.d), again with a more precise positioning of ADAP peptide 17-46 than peptide 97-126. In this binding mode, the ADAP-actin interaction should not interfere with barbed end binding, e.g. by formins or CapZ^23,24^. An overlay of the docked ADAP-actin structure with the CapZ/twinfilin/actin complex^25^ shows that simultaneous binding of all proteins is possible (Extended Fig. 8e). In support, in vitro actin polymerization by ADAP was enhanced several fold in the presence of CapZ (Fig. 4b). In contrast, superimposing the model structure of ADAP-actin with cofilin/actin^26^ shows that simultaneous binding is mutually exclusive (Extended Fig. 8f). Accordingly, in the presence of cofilin an initial increase in polymerization was followed by disassembly, likely reflecting cofilin-mediated F-actin depolymerization (Fig. 4c). Thus, ADAP synergises with CapZ, while it competes for the same or an overlapping site on filaments with cofilin.

## Discussion

Here, we show that the intrinsically disordered region of the scaffolding protein ADAP uses positively charged motifs to interact with G-actin. We propose a model where ADAP acts as a molecular sponge to locally enrich actin to a level above the critical concentration required for polymerization (Fig. 4d). Initial encounter of G-actin is possible at multiple sites in ADAP, leading to concentration enhancement of actin along a single ADAP molecule. Molecular crowding may further be enhanced by the concomitant interaction of one ADAP-bound actin monomer or dimer with a second ADAP molecule, which may itself be transiently bound to one or more actin molecules. When local concentrations of actin exceed the critical concentration, polymerization ensues. Once filaments are formed, ADAP remains bound to F-actin along the still available dimer interfaces.

This mode of action is distinct from the structured encounters that are a hallmark of actin regulators such as Arp2/3^27^, formins^23^ or WH2 domain containing proteins^28^. Given the colocalization of ADAP with key actin regulators, it is likely that the observed synergy *in vitro* (Fig. 4) is of functional relevance in the cellular environment. How the molecular crowding process described for ADAP is modulating actin dynamics in the context of the core regulators remains to be seen and relates to the question why only some cell types rely on the actin modulatory function of this protein. Speculatively, different cells and organisms have evolved different IDRs with the capacity to tune the self-assembly driven properties of the cytoskeleton in similar ways as suggested here for T cells.

## Materials and Methods

### Mice and isolation of T cells

ADAP-deficient mice were provided by Gary Koretzky^29^. WT and ADAP-deficient animals were under C57BL/6J background. All animal procedures were performed in accordance with the State of Sachsen-Anhalt, Germany and the Swiss legislation by the veterinary office of the Kanton of Bern, Switzerland. Splenic T cells were purified using T cell isolation kits and AutoMacs magnetic separation system (Miltenyi Biotec) and approximately 90% of the purified cells displayed a naïve phenotype as previously described^19^. 5C.C7 TCR^tg^ mice^30^ were maintained in pathogen-free animal facilities at the University of Bristol under the University mouse breeding Home Office License P10DC2972. All 5C.C7 TCR^tg^ mouse experiments had been approved by the University of Bristol AWERB (Animal welfare and ethical review body) committee.

### Live cell imaging using two-photon intravital microscopy

Isolated splenic wild type (WT) and ADAP-deficient T cells were fluorescently labeled with either CFSE or DDAO-SE (both from Invitrogen) and a total of 5×10^6^ cells were injected intravenously into the tail vein of WT recipient mice. Dyes were switched between experiments to exclude dye-related effects. After 2 h of adoptive transfer T cell interactions were imaged in the inguinal lymph node. Blood vessels were highlighted by additional injection of Rhodamine-Dextran/FITC-Dextran (1:1 v/v). Intravital imaging was performed as previously reported^31^. 2-Photon microscopy was performed using a ZeissLSM710 microscope equipped with a MaiTai DeepSee Femtosecond-Laser typically tuned to 800 nm on an AxioExaminer upright stage with a 20x, NA 1.0 water dipping lens. Image detection was done with three non-descanned (NDD) detectors typically equipped with emission detection filters of 565–610 nm (red), 500–500 nm (green), and ShortPass 485 nm (blue). Individual RGB scans in fields of view of typically 303×303 μm were recorded for time-lapse sequences typically every 4s and lasting up to 30 min. Image rendering was performed using Volocity 4.3 (Improvision, Waltham, MA, USA). Fluorescence-labeled T cells in contact with the vessel wall (visible by the rhodamin-dextran i.v. tracer) were indicated and contact time was estimated based on appearance in the individual image planes of the time lapse. Measured was the time before cells apparently present in the vessel lumen for at least two consecutive frames, i.e. at least 4 s, were floating away with the blood stream. A total of more than 100 T cells of each type in three independent experiments were quantified.

### Live cell imaging under shear flow

In vitro live cell imaging of T cell interaction behavior with endothelial cells under shear flow was performed as described before^32,33^. μ-dishes (Ibidi) were stimulated with TNF-α-(25 ng/mL; PromoKine) for 16-18 h. Before the experiments, pMBMECs were overlaid with 100 μM CCL21 (R&D systems) for 30 min at 37°C. Subsequently, the custom made flow chamber^32^ was mounted onto the pMBMECs and the μ-dish was placed onto the heated stage of an inverted microscope (Zeiss, AxioObserver). Flow was applied via an automated syringe pump (Harvard Apparatus. CMTMR- and blueCMAC-(both from Invitrogen)-labeled T cells in migration assay medium (MAM, DMEM with Glutamine (Gibco), 5% FCS, 25 mM HEPES) (20×10^6^ cells/mL of each, dyes were switched between experiments) were perfused over the pMBMECs at reduced flow of 0.25dyn/cm^2^ allowing for initial contact formation between the T cells and the pMBMECs. The dynamic T cell interaction was recorded under physiologic shear stress (1.5 dyn/cm^2^) with a monochrome CCD camera (AxioCam MRm Rev, Carl Zeiss) using a 20x objective (LD Plan Neofluar). Time-lapse videos were created from one frame every 20 sec over a recording time of 30 min.T cells were counted as arrested T cells if they attached on the endothelium during the accumulation phase (0.25 dyn/cm^2^) in the field of view (FOV) and resisted immediate detachment after one minute of physiological shear stress at 1.5 dyn/cm^2^. These arrested T cells were tracked manually using the ImageJ software (National Institute of Health) with manual tracking (Institute Curie, Orsay, France), chemotaxis, and a migration plugin (Ibidi). T cells that entered or left the FOV during the recording time were excluded from the analysis. The length/width ration for individual cells was calculated using the ImageJ software.

### F-actin content measurement by flow cytometry

Briefly, cells (0.1×10^6^ T cells in 100 μL PBS) were left untreated or were stimulated with anti-CD3 antibodies (145-2C11 (BD Bioscience) or OKT3 (eBioscience) each 10 μg/ml) or CXCL12/CCL21 (each 1 μg /ml R&D systems) for the indicated time points. The reaction was stopped by adding 100 μL FAT buffer (4% (v/v) paraformaldehyde (PFA), 1% (v/v) Triton-X containing FITC- or Alexa633-phalloidin (Sigma Aldrich/Invitrogen, 2 μg/mL) in PBS) and incubated for 15 min at room temperature. The cells were washed with PBS, resuspended in PBS/1% (v/v) PFA, and analyzed by flow cytometry (FACSCalibur, CellQuest™ Pro). The mean fluorescence intensity of untreated WT T cells or shC-transfected Jurkat T cells was set as 100%.

### Spreading assay

The spreading assay was performed as previously described^34^. Briefly, poly-L-lysine coated cover slides coated with anti-CD3 antibodies ((145-2C11 (BD Bioscience, 0.5 μg/μL) and OKT3 (eBioscience, 0.5 μg/μL)) or isotype controls (Biolegend, 0.5 μg/μL) were used to settle T cells at 37°C for 10 min. The reaction was stopped by washing the slide with ice-cold PBS. Cells were fixed, permealized and stained with rat mAb to ADAP (clone 92B2G10) in combination with anti-rat IgG-conjugated with FITC (Dianova). F-actin was stained with FITC- or TRITC-phalloidin (both from Sigma Aldrich). T cells were imaged on a LEICA TCS SP2 laser-scanning confocal system using 63x/1.32 oil objective. Multi-color overlays and quantification of the cell area were produced with Adobe Photoshop CS3 Extended software.

### Cell culture, transfection and Western Blotting

Jurkat T cells (ATCC) and B cells (Raji; ATCC) were maintained in RPMI 1640 medium (PAN) supplemented with 10% FBS (PAN) and stable L-glutamine at 37°C with 5% CO_2_. Jurkat T cells (2 × 107) were transfected by electroporation as previously described^15^. The knockdown efficiency of ADAP and SKAP55 was evaluated by Western blotting. Cell lysis were performed as previously described ^35,36^. Equivalent amounts of protein (50 μg of total protein) were separated by SDS-PAGE and transferred to nitrocellulose. Western blots were conducted with the indicated Abs and developed with the appropriate HRP-conjugated secondary Abs (Dianova) and the Luminol detection system (Carl Roth). The following antibodies were used for immunoblotting: anti-SKAP55 rat mAb (SK13B6^35^) anti-ADAP mAb (clone 5/FYB, BD Bioscience), anti-β-actin mAb (clone AC-15, Sigma-Aldrich). For quantification, the intensity of the detected bands was calculated using the Kodak Image station 2000R (ID image software).

### T cell interaction with B cells

Briefly Raji B cells were left untreated or were loaded with staphylococcal enterotoxin E (Toxin Technology, Inc.) and stained with DDAO-SE (red dye, Thermo Fischer Scientific). Raji B cells were incubated with an equal number of transfected Jurkat T cells (GFP-positive). Nonspecific aggregates were disrupted by vortexing, cells were fixed with 1% PFA (15 min, 4°C) and then analyzed by flow cytometry (FACSCalibur; CellQuest™ Pro). The percentage of conjugates was defined as the number of doublepositive (red and green signal) events. Imaging and image analysis protocols for the interaction of primary murine 5C.C7 T cells with CH27 B cell lymphoma APCs (RRID:CVCL_7178) have recently been described in great detail in a dedicated publication^37^. Briefly, time-lapse fluorescence microscopy was performed with primary 5C.C7 T cells transduced with recombinant Moloney Murine Leukaemia virus for the expression of fluorescent proteins, FACS-sorted to the lowest detectable sensor expression of 2 μM, and CH27 cells loaded with 10μM of the moth cytochrome C agonist peptide. All imaging was performed on a Perkin Elmer UltraVIEW ERS 6FE spinning disk confocal system fitted onto a Leica DM I6000 microscope equipped with full environmental control and a Hamamatsu C9100-50 EMCCD camera. A Leica 40x HCX PL APO oil objective (NA=1.25) was used for all imaging. Automated control of the microscope was performed with Volocity software (Perkin Elmer).

### Adhesion assay

For TCR-mediated adhesion 96-well plates were pre-coated with 0.5μg human ICAM-1/well (R&D systems). Transfected cells were left untreated or stimulated with anti-TCR antibody (OKT3, 5μg/ml) for 30 min at 37°C. Subsequently, cells were allowed to adhere for 30min at 37°C. Unbound cells were washed off with Hanks balanced salt solution (HBSS, PAN). Bound cells were counted and calculated as % input (2 × 10^5^ cells) in triplicates. For CXCR4-induced adhesion wells were coated with 0.5μg ICAM-1 together with CXCL12 (100ng/ml).

### Migration assay

Chemotaxis assays were performed using Transwells (6.5mm diameter insert, 5.0μm pore size, Costar) coated with human ICAM-1 (5μg/ml, R&D systems). Transfected Jurkat T cells (1 × 10^6^ cells/ml) were incubated for 2h at 37°C in the presence or absence of human CXCL12 (200ng/ml; lower chamber). The number of cells migrated into the lower chamber was counted and calculated as % input (of 0.5 × 10^6^ cells).

### Flow cytometric analysis of surface receptors

For cell surface expression on Jurkat T cells 0.2 × 10^6^ cells were incubated with anti-CD18 (clone MEM-48, antibodies-online.de), anti-CD184 (CXCR4, clone 12G5, BD Bioscience) or OKT3 (eBioscience) antibodies. After washing with PBS, bound antibodies on cells were detected with the APC-conjugated goat anti-mouse IgG (Dianova). Cells were analysed by flow cytometry (FACSCalibur, CellQuest™ Pro). Cells stained with the secondary antibody only were used as a reference.

### Construction of the suppression/re-expression vectors for ADAP deletion mutants

The generation of the suppression/re-expression vectors for ADAP has been previously described. The following primer pairs for the the cDNA constructs of ADAPΔ_1-340_ and ADAPΔ_1-381_ were used for cloning. ADAPΔ_1-340_: forward primer_340_Mlu1: CAACGCGTACCCCGAAACAGAAG, reverse primer_783_Not1: GGGCGGCCGCTTACTAGTCATTGTCATAGAT. ADAPΔ1-381 ADAPΔ1-340: forward primer_381_Mlu1: CAACGCGTAGCAAAGGCCAGAC reverse primer_783_Not1: GGGCGGCCGCTTACTAGTCATTGTCATAGAT PCR fragments were sequenced and cloned into the pCMS4 vector using the restriction sites MluI and NotI.

### Statistical analysis

Results are presented as means ± standard errors of the means (SEM). Unpaired Student’s t tests were used to assess the statistical significance using the GraphPad Prism software. A p ≤ 0.05 was considered statistically significant.

### Protein expression and purification

ADAP-full length and all the ADAP-fragments were cloned in pET28a vector and transformed in BL21 (rosetta) competent cells for protein expression. These proteins were generated under the expression of T7 promoter in complex medium (2YT) for normal experiments and minimal (M9) medium for NMR experiments. All the proteins were N-terminally fused to a 6x histidine-tag, purified by Ni-NTA affinity chromatography using 250mM imidazole in the elution buffer. To co-express ADAP and SKAP55 proteins in pFastBacDual vector, insect cell (Sf9) expression system was used. The complex was purified through Strep-tag affinity purification using 5mM desthiobiotin or 50mM biotin in the elution buffer. All the proteins after affinity purification were further purified by size exclusion chromatography on a superdex_200 increase 10/300 gl column in a final buffer containing 20mM Hepes, and 50mM NaCl of pH 7.5. The human β-actin used in the NMR experiments was recombinantly expressed using Sf9 insect cells (as described in ^38^). The construct was kindly provided by Shimada group from University of Tokyo. The non-polymerizable monomeric actin was bought from Hypermol company.

### *In vitro* polymerization assay

Pyrene-labeled monomeric actin (G-actin) from rabbit skeletal muscle was used for these assays (purchased from Hypermol company). Different concentrations of test proteins were added to 5μM of pyrene actin and change in fluorescence intensity was measured over time at an interval of 10 seconds on a TECAN plate reader. The protein buffer mixed with pyrene actin was used as a control to rule out the possibility of spontaneous polymerization induced by buffer in which proteins are eluted and stored. The excitation wavelength was 385nm and emission was measured at 420nm ± 10nm. The data points were plotted as fluorescence intensity against time (Origin_2019b 64bit software). The half time to maximal polymerization was calculated as described in^39^.

### F-actin bundling / co-sedimentation assay

The monomeric actin was polymerized with actin polymerization buffer (also called F-buffer containing 1M KCl and 10mM ATP, pH 8) to a concentration of 10μM and incubated at room temperature for 15-30 minutes to ensure the complete polymerization. Then 10μl of 10μM or 20μM test protein was added to the prepolymerized actin and again the mixture was incubated for 30 minutes followed by centrifugation for 15 minutes at 12000g at RT. Several controls were used to validate the results such as a control without F-actin (i.e., test protein with F-buffer) to rule out the possibility of self-precipitation. Furthermore, F-actin in protein buffer was used to make sure that F-actin itself does not form bundles. BSA was utilized as a negative control and α-actinin as a positive control. After the centrifugation step, the supernatants were carefully removed to a new tube and the invisible pellet was resuspended in 10μl water. The supernatants and pellets were ultimately run on SDS PAGE and analyzed using Image J software. The plots and error bars were calculated on GraphPad prism software.

### NMR sample preparation and measurement

For the interaction experiments with actin, ADAP proteins were recombinantly expressed under deuterated conditions containing ^15^N-NH_4_Cl and deuterated glucose. ADAP_1-100_ was expressed in non-deuterated conditions. The overexpressed proteins were purified as described above. ^1^H-^15^N-HSQC spectra were acquired on a Bruker Avance III 700 MHz spectrometer equipped with a 5 mm triple resonance cryoprobe at 300K with 32 scans (deuterated) or 64 scans (non-deuterated) and 160 data points in the indirect dimension. 25μM ^2^H-^15^N- or ^15^N-labeled protein was mixed with 25μM G-actin (recombinantly expressed) or non-polymerizing actin (purchased from Hypermol company) in a final volume of 550μl containing 10% D_2_O and 10mM HEPES, 50mM NaCl, 0.4mM ATP, pH 6.5. NMR spectra were processed with TopSpin3.2 (Bruker, Billerica, USA) and analyzed using CcpNMR Analysis (version 2.4.2.)^40^.

For backbone assignments ^13^C-^15^N-labeled ADAP_1-100_ and ADAP_100-200_ were recombinantly expressed in M9 minimal medium containing ^15^N-NH_4_Cl and ^13^C-glucose. The overexpressed proteins were purified as described above and measurements were performed in 10mM HEPES, 50mM NaCl, pH 6.5 with 10% D2O. HNCA, HN(CO)CA, HNCO, HN(CA)CO, HNCACB spectra were recorded at 300K for 500μM ^13^C-^15^N-labeled ADAP_1-100_. HNCA, HN(CO)CA, HNCO, HN(CA)CO, HNCACB, (H)N(CA)NH spectra were recorded at 300K and HNCA, HN(CO)CA, (H)N(CA)NH were recorded at 280K for 400μM ^13^C-^15^N-labeled ADAP_100-200_. 3D experiments were acquired with 8-32 scans and 25-50% NUS. Backbone assignments for 85% of the non-proline residues of ADAP1-100 (at 300K, BMRB ID 51216) and 75% of the non-proline residues of ADAP100-200 (at 280K, BMRB ID 51217) could be assigned using CcpNMR Analysis (version 2.4.2.)^40^. Overlapping peaks were excluded from the peak intensity analysis. Chemical shift differences were calculated using the formula Δδ(^1^H,^15^N)=(δ(^1^H)^2^+(0.15*δ(^15^N))^2^)^1/2^.

### Electron microscopy

Samples from in vitro polymerization assay were diluted to 50-100 ng/μl and 4 μl were applied to plasma-treated formvar/carbon-coated copper grids (S162-4, Plano GmbH, Wetzlar, Germany) for 1 minute. Grids were blotted manually with Whatman No.1 filter paper and stained with 1% uranyl acetate for 1 minute. After blotting, grids were dried and stored at room temperature until use. Imaging was conducted on a FEI Talos L120C TEM operated at 120 kV equipped with a CETA 16M detector under low dose conditions using TIA software (FEI company, Eindhoven). Exposures were taken at a nominal defocus of −1.5 μm. Analysis of filament diameters was conducted manually with ImageJ. In short, the “straight” tool was used to create selections of twice the diameter of the filament using a line width of the filament diameter. Lines were arranged perpendicularly to the filament and used to plot a density profile for reading off the diameter.

### Crosslinking-mass spectrometry sample preparation

5μM of non-polymerizing actin or pre-polymerized actin were mixed with 5μM of ADAP-full-length or different fragments of ADAP and cross-linked with1mM DSSO for 20 minutes at room temperature and quenched with 20mM Tris-HCl pH 8.0. As described previously^41^, crosslinked proteins were denatured and reduced using crystallized urea and DTT to a final concentration of 8M and 1mM, respectively. The proteins were further lysed by trypsin and the peptides were desalted using C18 stage-tips and dried in vacuum centrifuge.

### Measurement and analysis of DSSO cross-linked samples by mass spectrometry

The dried peptides were dissolved in 100 μl 1% ACN / 0.05 % TFA and injection volume of 1 μl was used. LC-MS analysis was performed using an UltiMate 3000 RSLC nano LC system coupled on-line to an Orbitrap Fusion mass spectrometer (Thermo Fisher Scientific). Reversed-phase separation was performed using a 50 cm analytical column (in-house packed with Poroshell 120 EC-C18, 2.7μm, Agilent Technologies) with a 120 min gradient. Cross-link acquisition was performed using an LC-MS2 method. The following parameters were applied: MS resolution 120,000; MS2 resolution 60,000; charge state 4-8 enabled for MS2; stepped HCD energy 21, 27, 33. Cross-linking Data analysis was performed using XlinkX standalone^41^ with the following parameters: minimum peptide length=6; maximal peptide length=35; missed cleavages=3; fix modification: Cys carbamidomethyl=57.021 Da; variable modification: Met oxidation=15.995 Da; DSSO cross-linker=158.0038 Da (short arm = 54.0106 Da, long arm = 85.9824 Da); precursor mass tolerance = 10 ppm; fragment mass tolerance = 20 ppm. Database search was performed using the protein sequences of actin and ADAP.

### Peptide docking models

The CABS-dock server (http://biocomp.chem.uw.edu.pl/CABSdock)^42,43^ was used to dock fragments of ADAP_1-381_ to monomeric actin (PDB 3hbt)^44^ or an actin dimer (chains C and E extracted from PDB 6fhl^45^). A soft distance restraint of 20Å between Lysine residues was included in the coarse grained model.

## Supporting information

Supplementary_Info_Movie_References

## Authors contribution

C.F., J.S. and N.D. designed and interpreted all experiments for the structural, biophysical and biochemical analysis. S.K. and B.S. designed the experiments using the ADAP KO mice. P.R. and J.D performed and analyzed the 2-Photon microscopy studies. J.D., R.L, M.A. and M.H. performed shear flow experiments and analyzed the data. J.D. and S.K. performed confocal microscopy studies and determined the F-Actin content in response to chemokine and TCR triggering in T cells. S.K. and C.M. performed and analyzed the functional assays using Jurkat T cells. N.D. performed protein purification, polymerization, bundling experiments and prepared samples for NMR, crosslinking MS and negative stain EM. J.S. performed and analyzed NMR experiments and performed the molecular docking. F.L. and H.S. performed crosslinking mass spectrometry experiments. T.W. contributed to the cloning of constructs. B.K. performed quality control experiments of the ADAP-actin samples by mass spectrometry and helped with the analysis of cross-linking-MS data. T.H. performed EM experiments and analyzed the data. C.W. performed live cell microscopy experiments with ADAP and SKAP55. C.F. and S.K. and B.S. supervised the project. C.F. wrote the manuscript with contributions from all authors.

## Acknowledgement

We thank A. Ramonat, G. Höbbel, L. Philipsen and J. Dudeck for excellent technical assistance and Dr. Miguel Álvaro-Benito for helpful discussions. We are also thankful to Jens V. Stein (Department of Oncology, Microbiology and Immunology, University of Fribourg, Fribourg, Switzerland) who took care about the animal procedure for the Swiss legislation by the veterinary office. This work was supported by the Deutsche Forschungsgemeinschaft grant CRC854 project B12 (S.K. and C.F.). We would like to acknowledge the assistance of the Core Facility BioSupraMol supported by the DFG for NMR, MS and cryoEM measurements.

