## Extended data Figures for "ADAP’s intrinsically disordered region is an actin sponge regulating T cell motility"

Extended Data Fig. 1

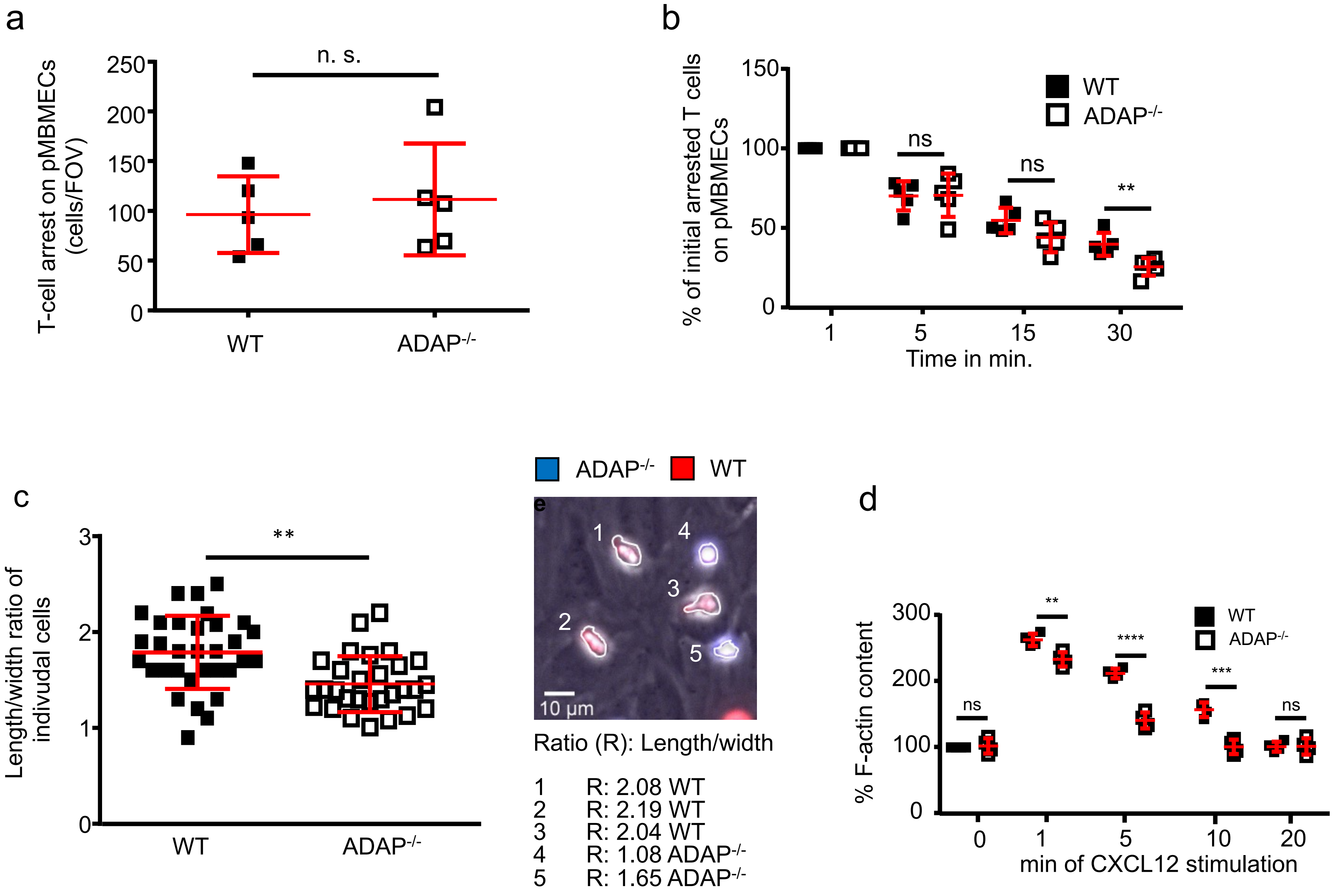

Extended Data Fig. 1

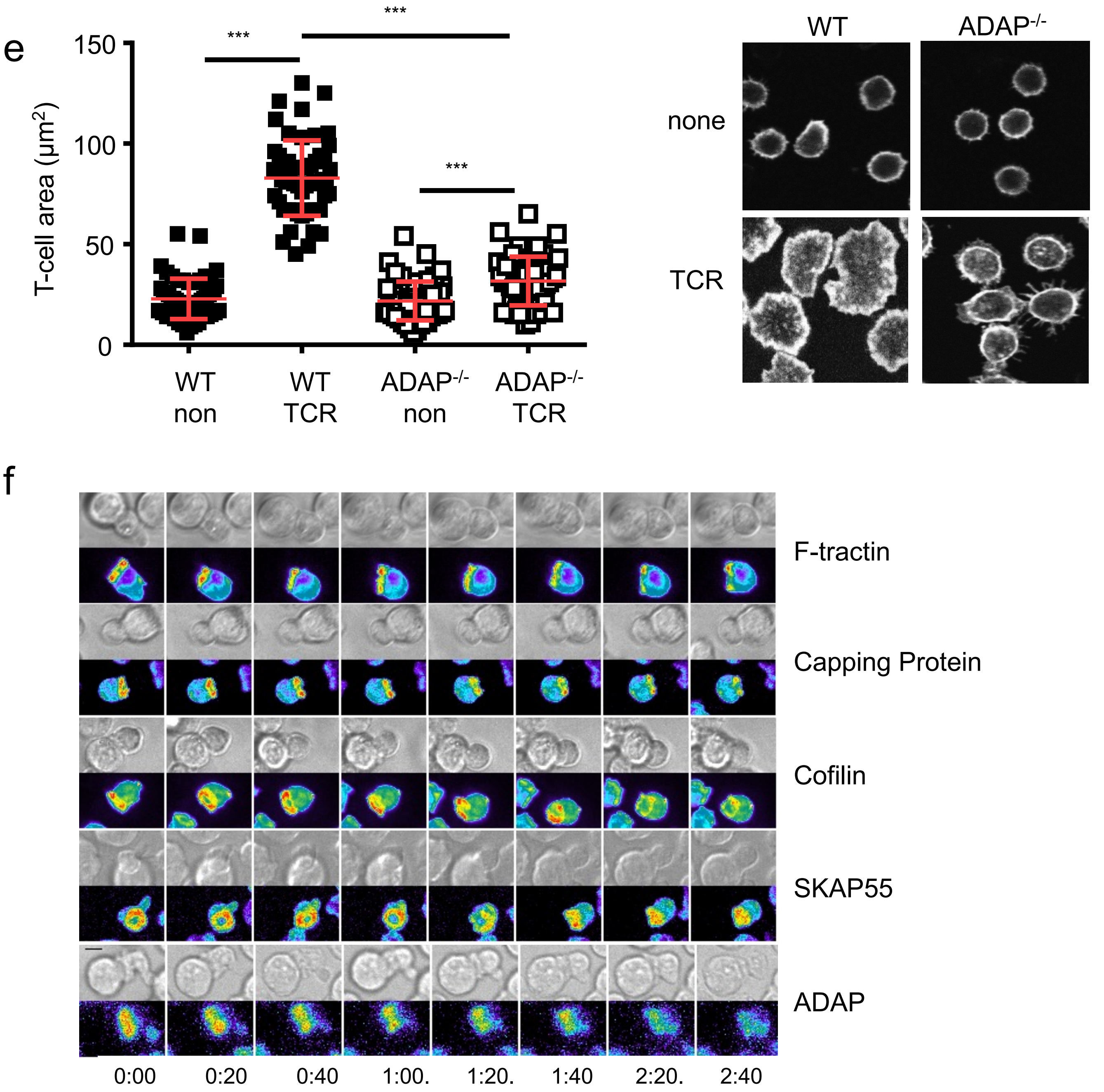

**Extended figure 1: Loss of ADAP affects stable arrest under shear flow, polarization, CXCR4-induced F-actin content and TCR-mediated spreading in T cells *ex vivo*.** (a-d) Purified fluorescence-labeled WT and ADAP$^{-/-}$ T cells were perfused in a parallel flow chamber. Low shear stress (0.25 dyn/cm$^2$) allowed T cells to accumulate on a monolayer of CCL21-coated TNF-$\alpha$-stimulated pMBMECs. After 2 min, the shear stress was increased-to-physiological strength (1.5 dyn/cm$^2$), and T-cell interactions with pMBMECs were recorded under constant flow for 30 min. (a) Shear-resistant arrested WT and ADAP$^{-/-}$ T cells were counted after 1 min of increased to physiological shear per field of view (FOV). (b) The remaining WT and ADAP$^{-/-}$ T cells were represented as a percentage of initially arrested WT T cells or ADAP$^{-/-}$ T cells on pMBMECs after 5, 15, and 30 min of physiological shear stress (n=2, 5 films, **p<0.01). (c) The length/width ratio was used to determine the polarization status of individual cells after 10 min of physiological shear (n=30-40 cells, 2 films, **p<0.01). (d) Representative image of polarized WT (red) and non-polarized ADAP-/- (blue) T cells on CCL21-coated TNF-α-stimulated pMBMECs during shear flow. The length/width ratio of individual cells is depicted. (e) Isolated WT and ADAP$^{-/-}$ T cells were left untreated or stimulated with CXCL12 for the indicated time points. T cells were permeabilized, fixed, and stained with FITC-Phalloidin. The F-actin content was measured by flow cytometry. The mean fluorescence intensity of untreated WT T cells was set to 100% (n=4, **p<0.01, ***p<0.001, ****p<0.0001). (f) Purified WT and ADAP$^{-/-}$ T cells were plated on isotype coated (non) or anti-CD3 mAb (TCR)-coated poly-L-lysine slides. T cells were fixed, permeabilized, and stained with FITC-Phalloidin to visualize the F-actin. T cells were imaged by confocal microscopy to determine the T-cell area (n=3, 60 WT and ADAP$^{-/-}$ T cells, ***p<0.001). Representative T cells are depicted. (g) ADAP localization is comparable to that of key actin regulators. 5C.C7 TCR transgenic primary mouse T cells were retrovirally transduced to express the indicated protein tagged with GFP. Transduced T cells were activated with CH27 B cell lymphoma antigen presenting cells incubated with the moth cytochrome C 89-103 peptide recognized by the 5C.C7 TCR. Duration after tight cell coupling is given with 0:00 the time of tight cell couple formation. Representative imaging data are given with a differential interference contrast (DIC) image on top and the maximum projection of the three-dimensional GFP fluorescence data at the bottom in a rainbow false color intensity scale. Black scale bar=5µm. Subcellular distributions of core actin regulators are shown to be similar by computational quantification in Roybal et al. (2016)[46]. Extension of such analysis to ADAP was prevented by limiting retroviral transduction efficiency.

Extended Data Fig. 2

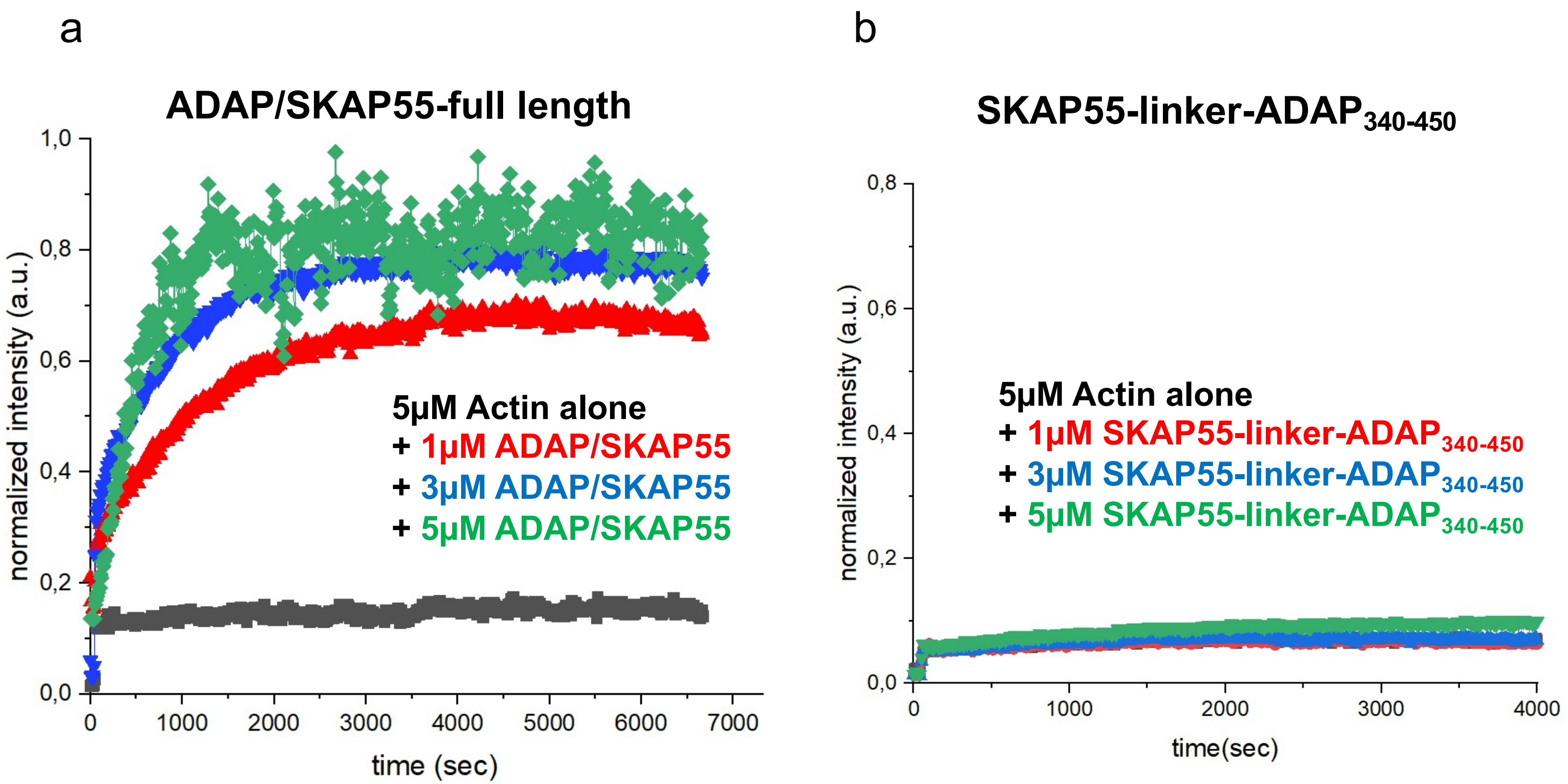

### Extended data Fig. 2

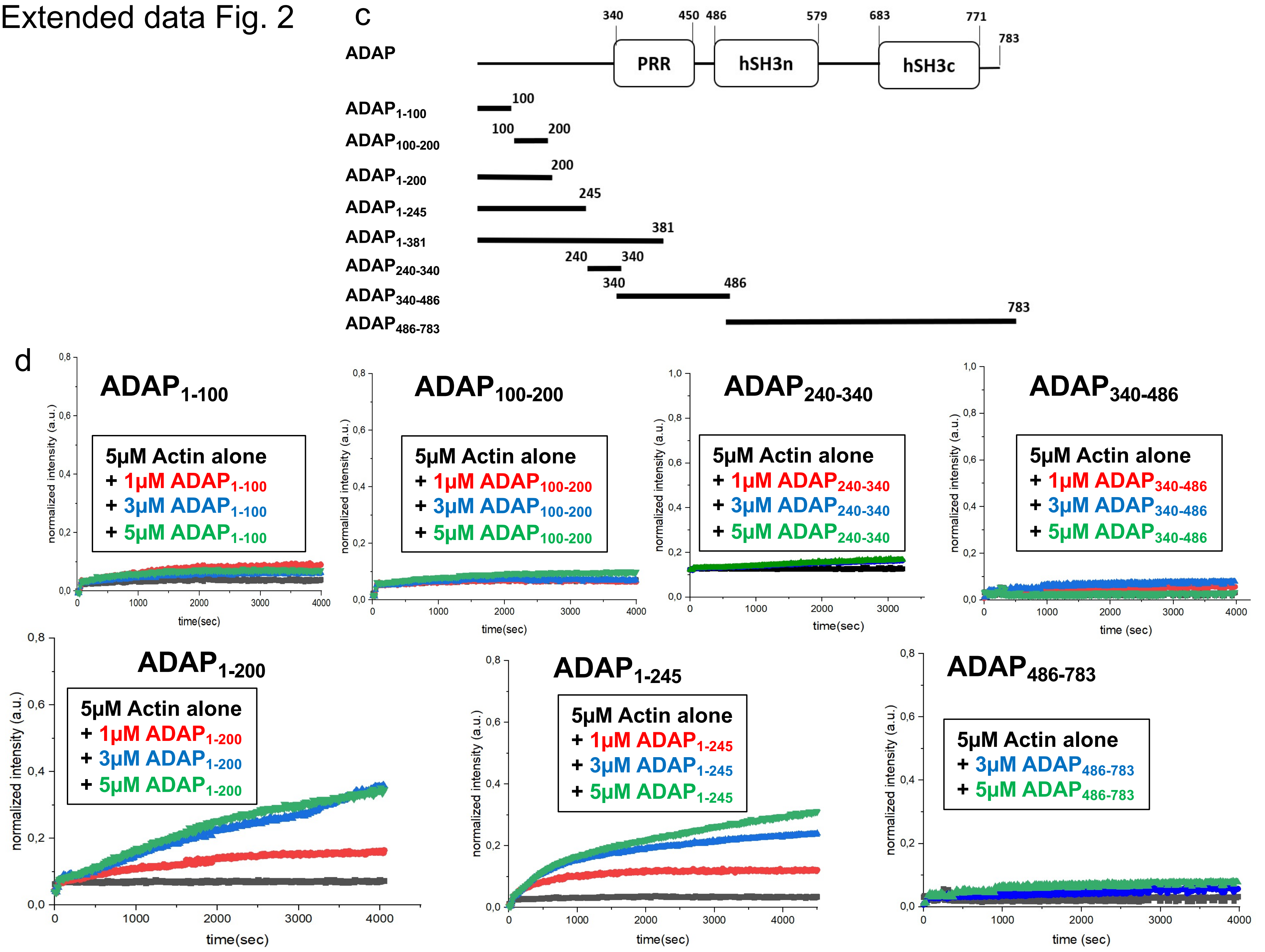

**Extended Data Fig. 2: *In vitro* actin polymerization by ADAP/SKAP55 complex and different fragments of ADAP.** (a) Polymerization curves representing the change in fluorescence intensity of pyrene actin with respect to time at different concentrations (1μM, 3μM and 5μM) of ADAP/SKAP55 complex. (b) SKAP55 protein linked to $ADAP_{340\text{-}450}$ showed no polymerization activity. (c) Various fragments of ADAP generated to narrow down the region responsible for the polymerization activity. All the fragments were cloned in pET28a vector with an N-terminal His-tag and expressed in bacterial cell expression system. The over-expressed proteins were then purified by affinity tag purification and re-purified by size exclusion chromatography and were checked for their polymerization ability. The constructs comprising a length of 100-150 amino acids such as 1-100, 100-200, 240-340 and 340-486 showed no significant polymerization activity. The pyrene-actin concentration of 5μM was kept constant in all the samples, change in fluorescence intensity was measured on TECAN plate reader at an interval of 10 seconds.

### Extended figure 3

ADAP$_{1-381}$

a

```
MAKYNTGGNP TEDVSVNSRP FRVTGPNSSS GIQARKNLFN NQGNASPPAG
        60         70         80         90        100
PSNVPKFGSP KPPVAVKPSS EEKPDKEPKP PFLKPTGAGQ RFGTPASLTT
       110        120        130        140        150
RDPEAKVGFL KPVGPKPINL PKEDSKPTFP WPPGNKPSLH SVNQDHDLKP
       160        170        180        190        200
LGPKSGPTPP TSENEQKQAF PKLTGVKGKF MSASQDLEPK PLFPKPAFGQ
       210        220        230        240        250
KPPLSTENSH EDESPMKNVS SSKGSPAPLG VRSKSGPLKP AREDSENKDH
       260        270        280        290        300
AGEISSLPFP GVVLKPAASR GGPGLSKNGE EKKEDRKIDA AKNTFQSKIN
       310        320        330        340        350
QEELASGTPP ARFPKAPSKL TVGGPWGQSQ EKEKGDKNSA TPKQKPLPPL
       360        370        380
FTLGPPPPKP NRPPNVDLTK FHKTSSGNST S
```

Deletion 1
Deletion 2
Deletion 3
Deletion 4
Deletion 5
Deletion 6
Deletion 7
Deletion 8

b

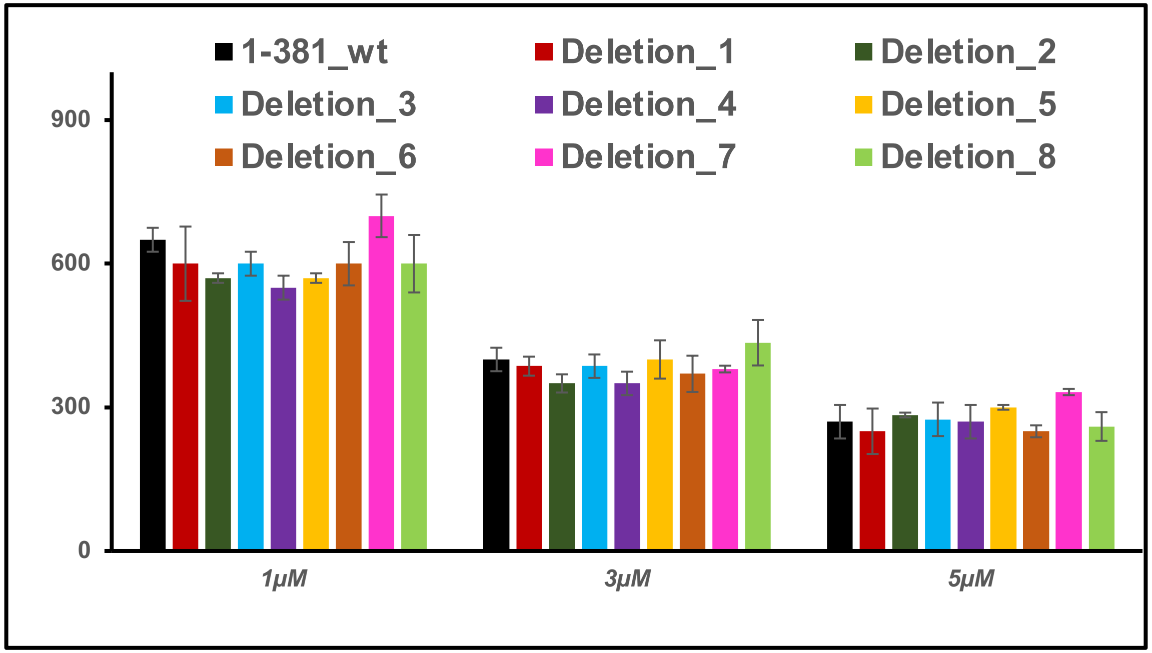

**Extended Data Fig. 3: Deletions in $ADAP_{1-381}$ do not impair ADAP's ability to polymerize actin.** (a) Series of deletion mutants named 1-8 were created in the sequence of $ADAP_{1-381}$. (b) The impact of each deletion was checked on actin polymerization and further compared to the activity of wild-type $ADAP_{1-381}$ (Fig. 2b). All the deletion constructs showed activity similar to wild-type $ADAP_{1-381}$ when used in *in vitro* polymerization assay.

### Extended Data Fig. 4

a

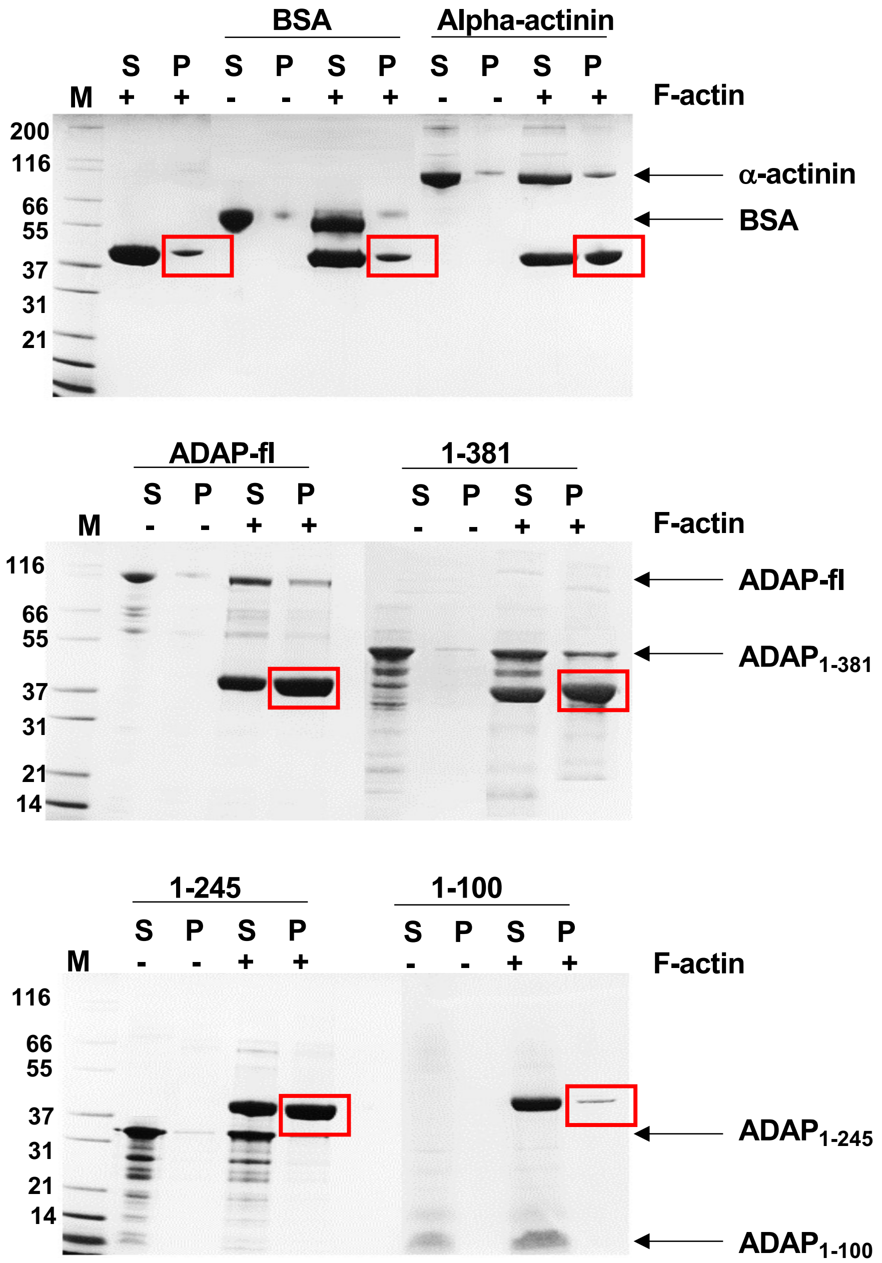

### Extended Data Fig. 4

b c d

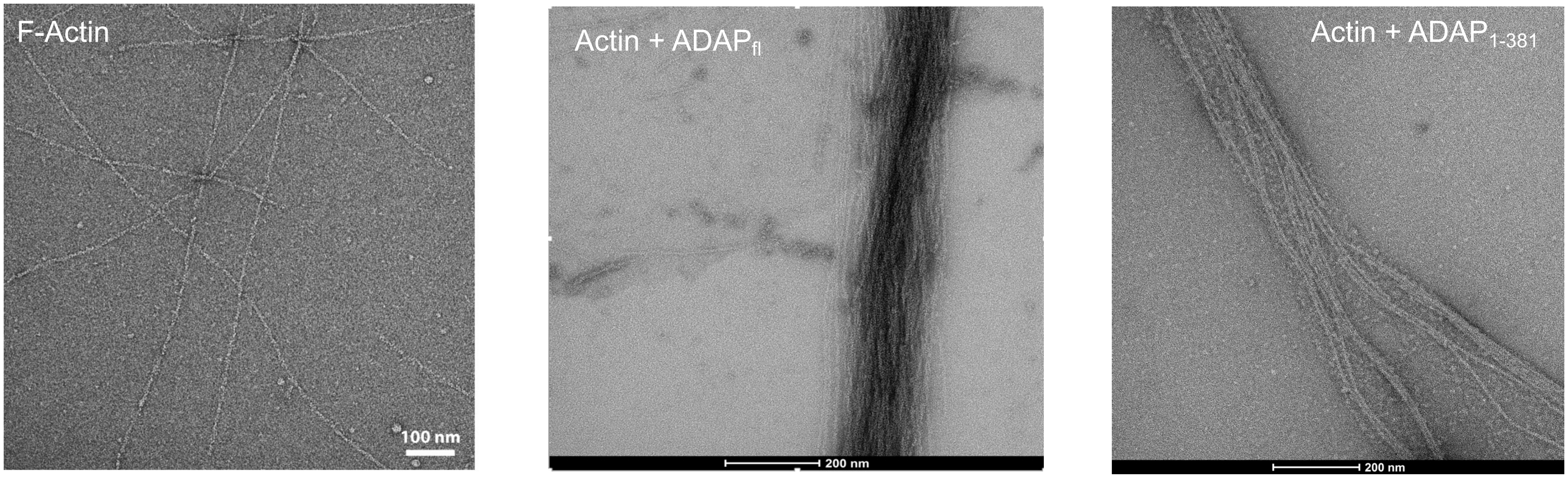


e

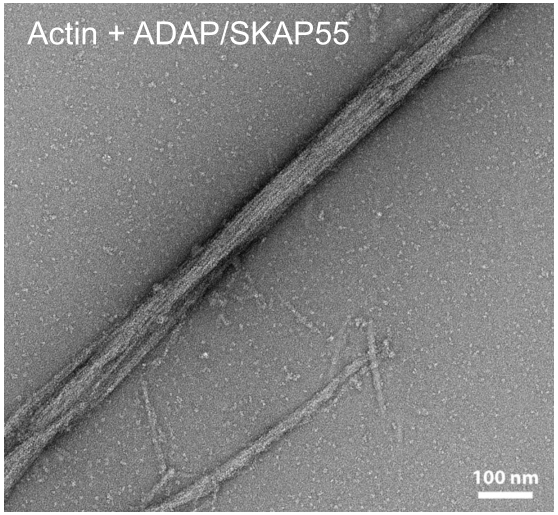


f

| Sample | min diameter (nm) | max diameter (nm) | mean (nm) | std.dev. |
|---|---|---|---|---|
| F-Actin | 5 | 8.6 | 6.9 | 1.03 |
| ADAP-SKAP55 | 50 | 75 | 63 | 7.01 |
| ADAP | 8 | 800 | n.d. | n.d. |
| ADAP1-381 | 14 | 500 | n.d. | n.d. |

**Extended Data Fig. 4: ADAP induces bundling of actin filaments *in vitro*.** (a) SDS-PAGE analysis showing the bundling of F-actin by different ADAP constructs. F-actin was mixed with different ADAP fragments and centrifuged at low speed for 30 minutes at room temperature. The supernatant and the resuspended pellet were analyzed on SDS PAGE. ADAP-full length and the N-terminal ADAP-fragments (1-245, 1-381), similar to the positive control ($\alpha$-actinin), show bundling of filamentous actin (F-actin) as compared to the negative controls (BSA or F-actin alone). Arrows highlight the respective ADAP constructs, and red rectangles indicate sedimented actin. (b-e) Negative stain EM of actin filaments (b) and bundles formed upon addition of (c) ADAP-full length, (d) $ADAP_{1\text{-}381}$ and e) ADAP/SKAP55 complex. f) shows the summary of analysis for several negative stain micrographs for the constructs shown in b)-e)

Extended Data Fig. 5

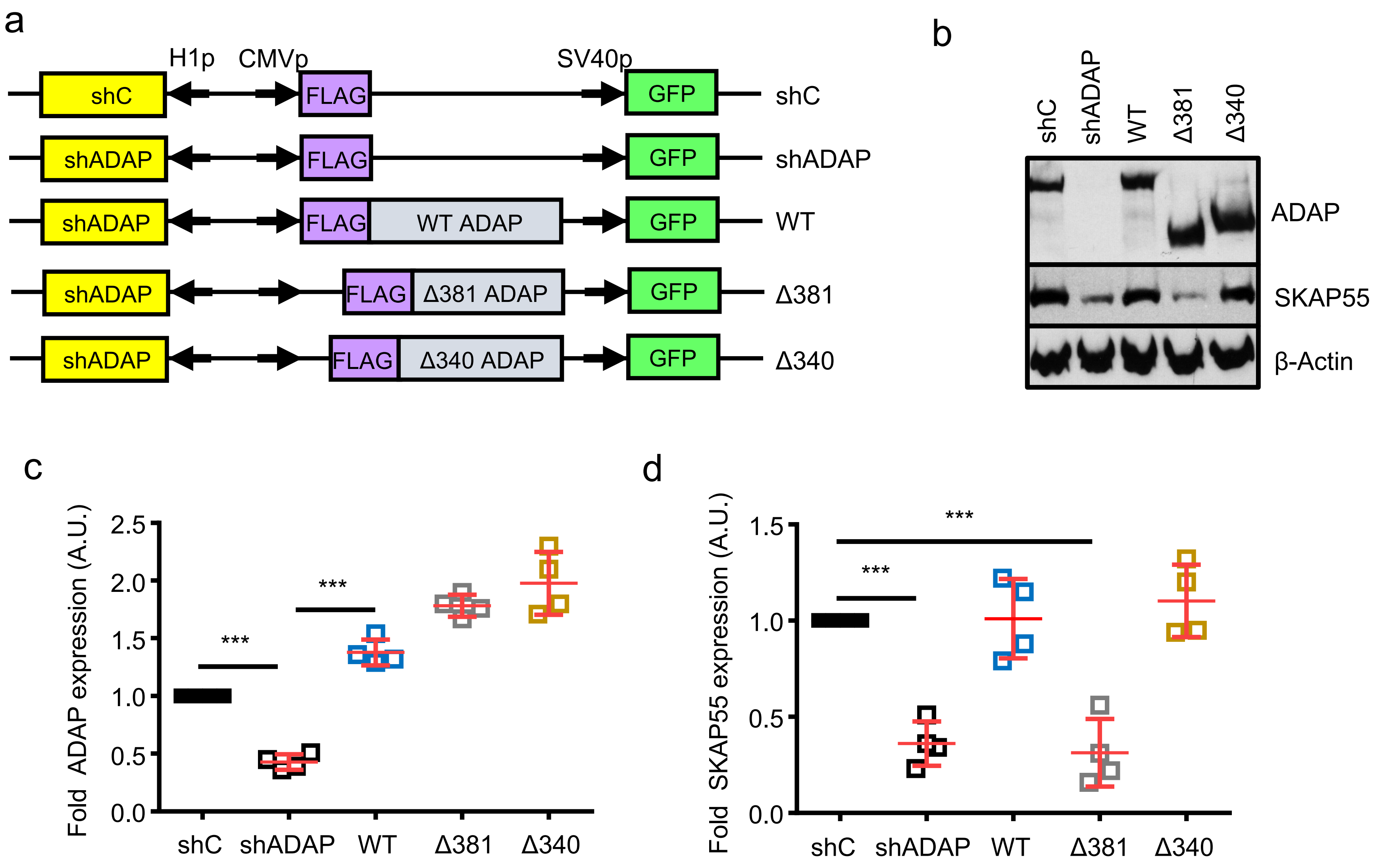

Extended Data Fig. 5

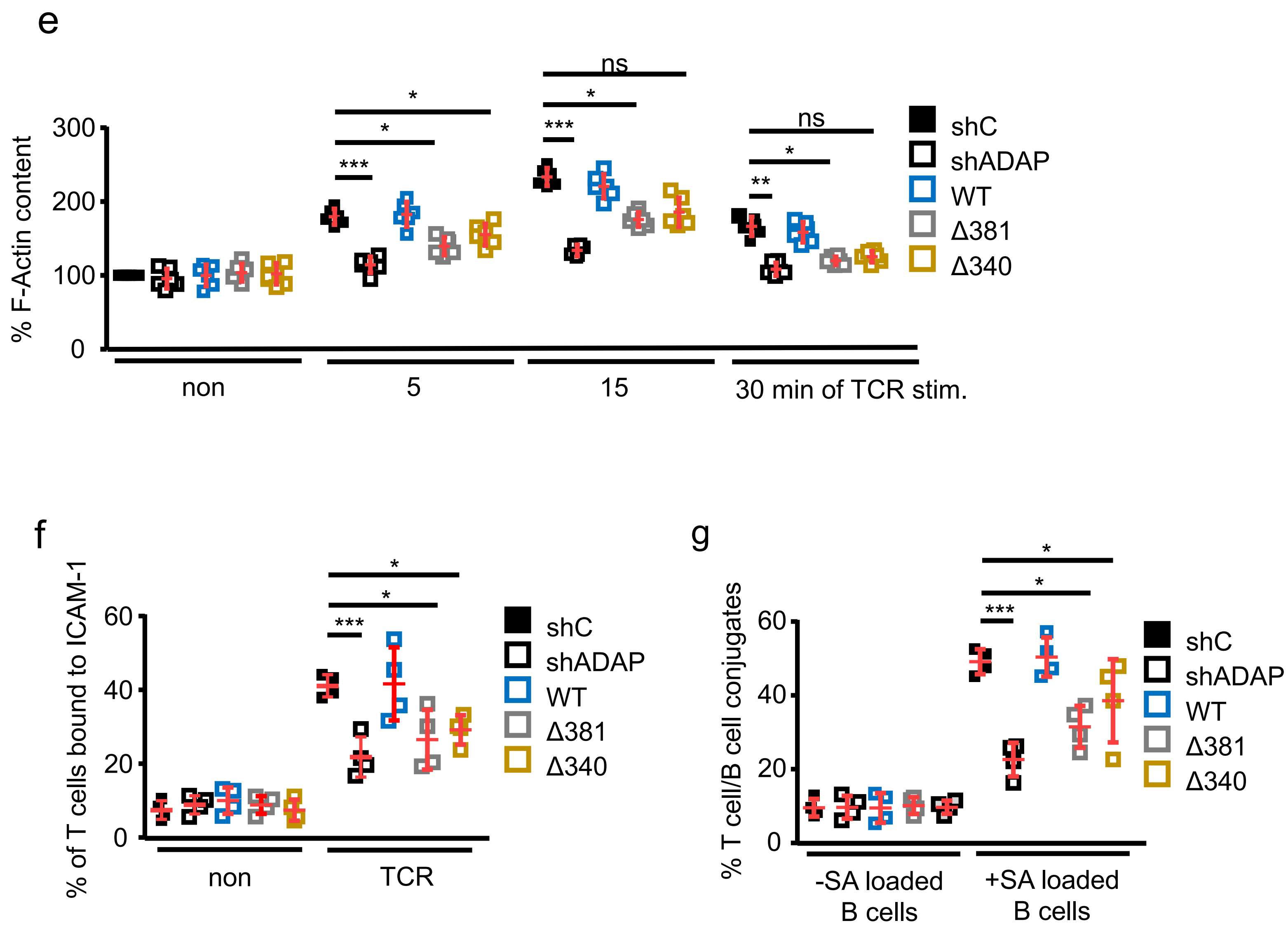

### Extended Data Fig. 5

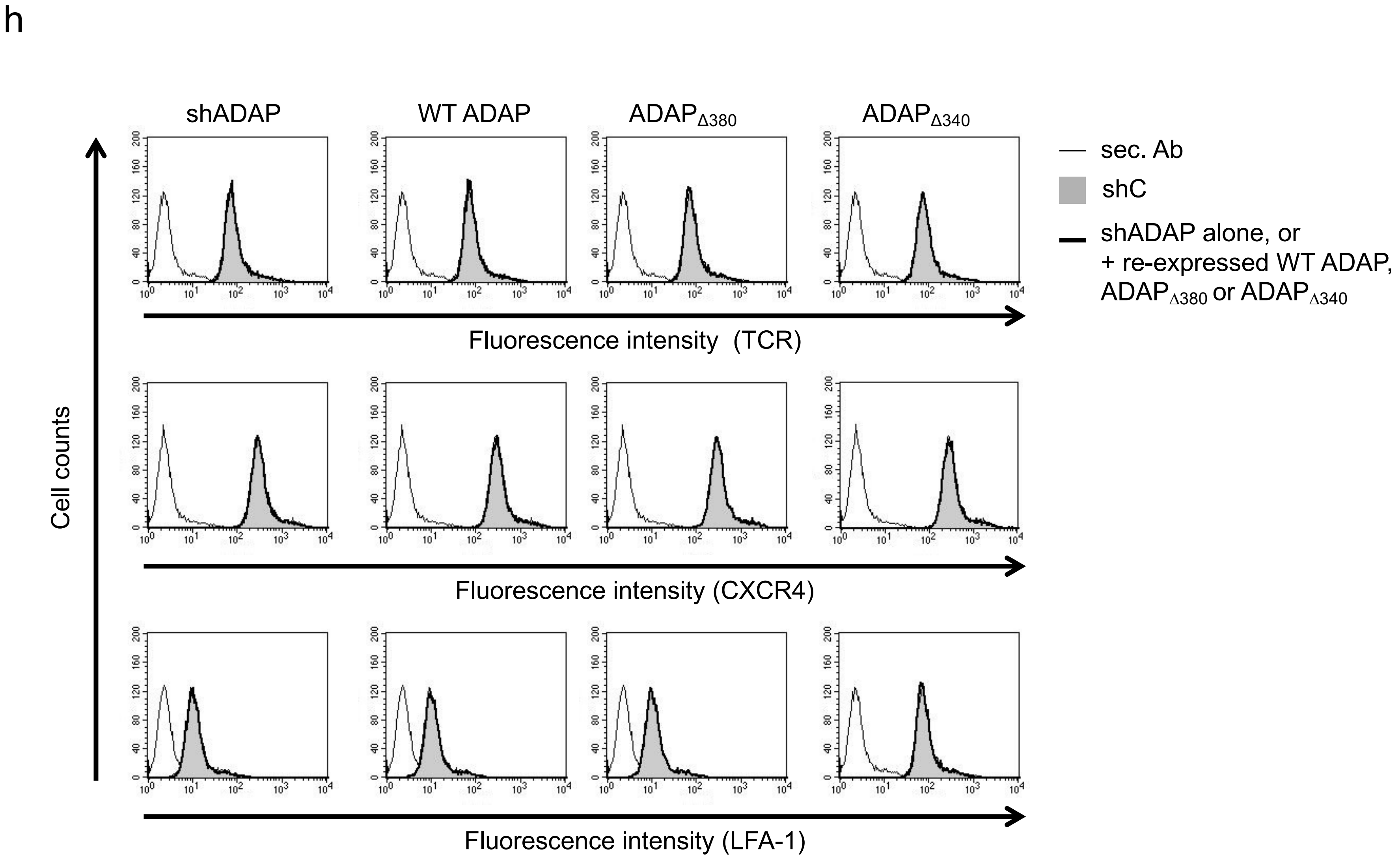

**Extended figure 5 : Deletion of the N-terminus of ADAP controls TCR-mediated F-actin content, adhesion and interaction of T cells with APCs.** (a) Schematic representation of the suppression/re-expression plasmids for ADAP used in this study. (b) Jurkat T cells were transfected with suppression/re-expression plasmids which do not suppress endogenous ADAP (shC), reduce the endogenous protein level of ADAP (shADAP), re-express a FLAG-tagged shRNA-resistant wild type ADAP (WT) or re-express two N-terminal deletion mutants of ADAP (Δ340 and Δ381). 48 hr after transfection, lysates were analyzed by Western blotting for ADAP, SKAP55 and β-actin (loading control). (c-d) Quantification of the reduction and re-expression of ADAP and its mutants after normalization to the ADAP and SKAP55 expression levels of the shC-tranfected control cells, which were set to 1 (c for ADAP and d for SKAP55). (e) After transfected T cells were stimulated with anti-CD3 antibodies for the indicated times points, permeabilized, fixed, and stained with Alexa633-Phalloidin to visualize F-actin. The amount of F-actin in GFP-expressing cells was determined by flow cytometry. The F-actin content is shown as the percentage of F-actin in shC-transfected untreated T cells, which was set to 100 %.(f) Transfected Jurkat T cells were analyzed for their ability to adhere to ICAM-1-coated wells in a resting state or stimulated for 30 min with anti-CD3 mAbs. Adherent cells were counted and calculated as percentage of input (n=4). g) Cells were transfected as described in (b) and analyzed for their ability to form conjugates with DDAO-SE (red)-stained Raji B cells that were pulsed without (non) or with superantigen (SA) for 30 min. The percentage of conjugates was defined as the number of double positive events in the upper right quadrant (n=4,$^*p<0.05$, $^{**}p<0.01$, $^{***}p<0.001$). h) Expression levels of the TCR, CXCR4 and CD18 of transfected Jurkat T cells. Jurkat T cells were transfected with suppression/re-expression constructs that do not suppress endogenous ADAP (shC) or reduces the protein level of ADAP (shADAP) and re-express a FLAG-tagged shRNA-resistant wild type form of ADAP (WT ADAP) or its deletion mutants ($ADAP_{\Delta 380}$ and $ADAP_{\Delta 340}$). Transfectants were used for flow cytometric analysis of the TCR, CXCR4 and CD18 surface expression within the GFP gate (n=3). One representative experiment out of three is shown.

Extended Data Fig. 6

a

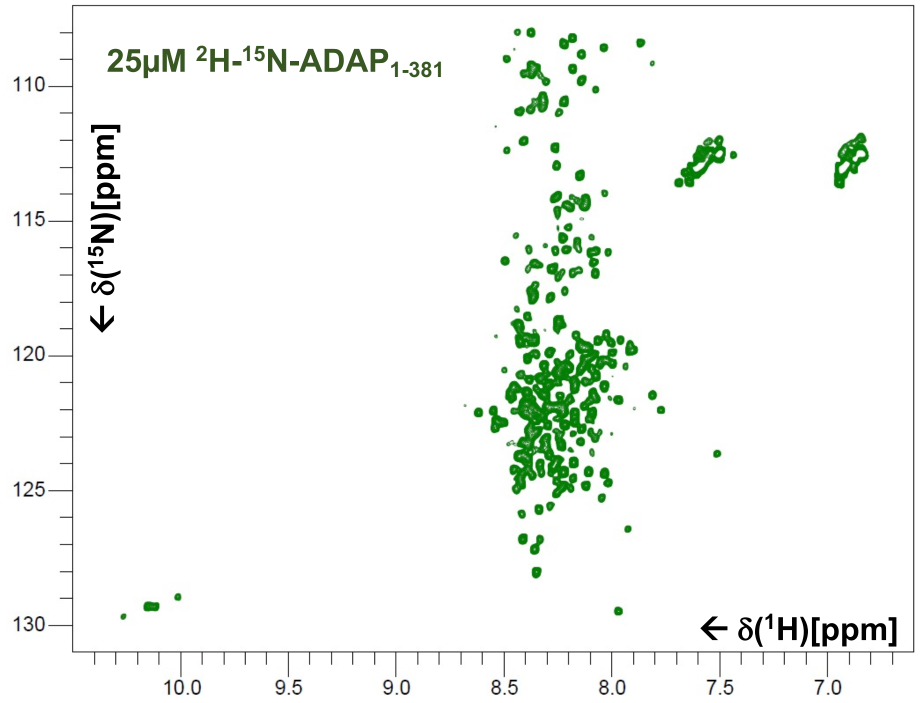


b

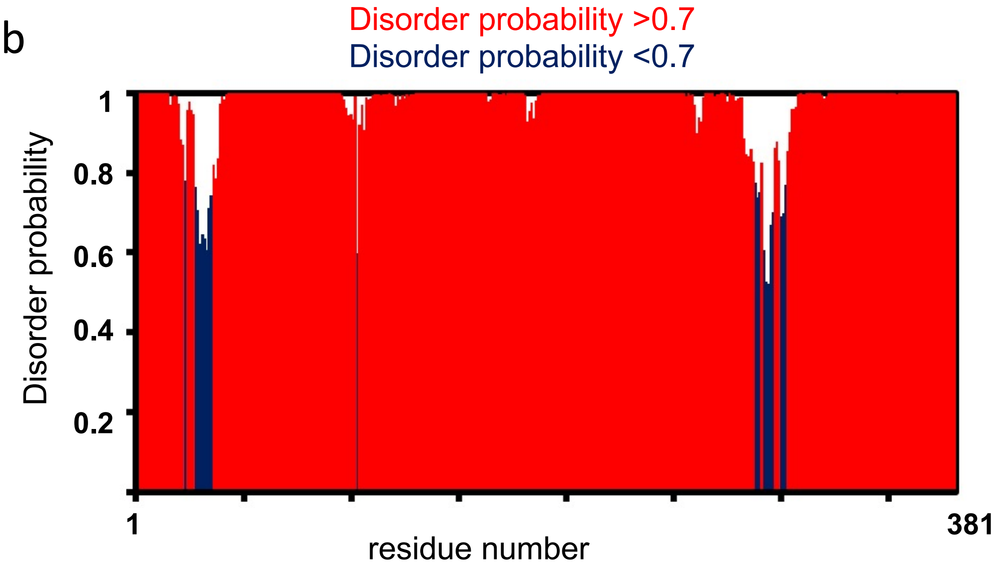

Extended Data Fig. 6

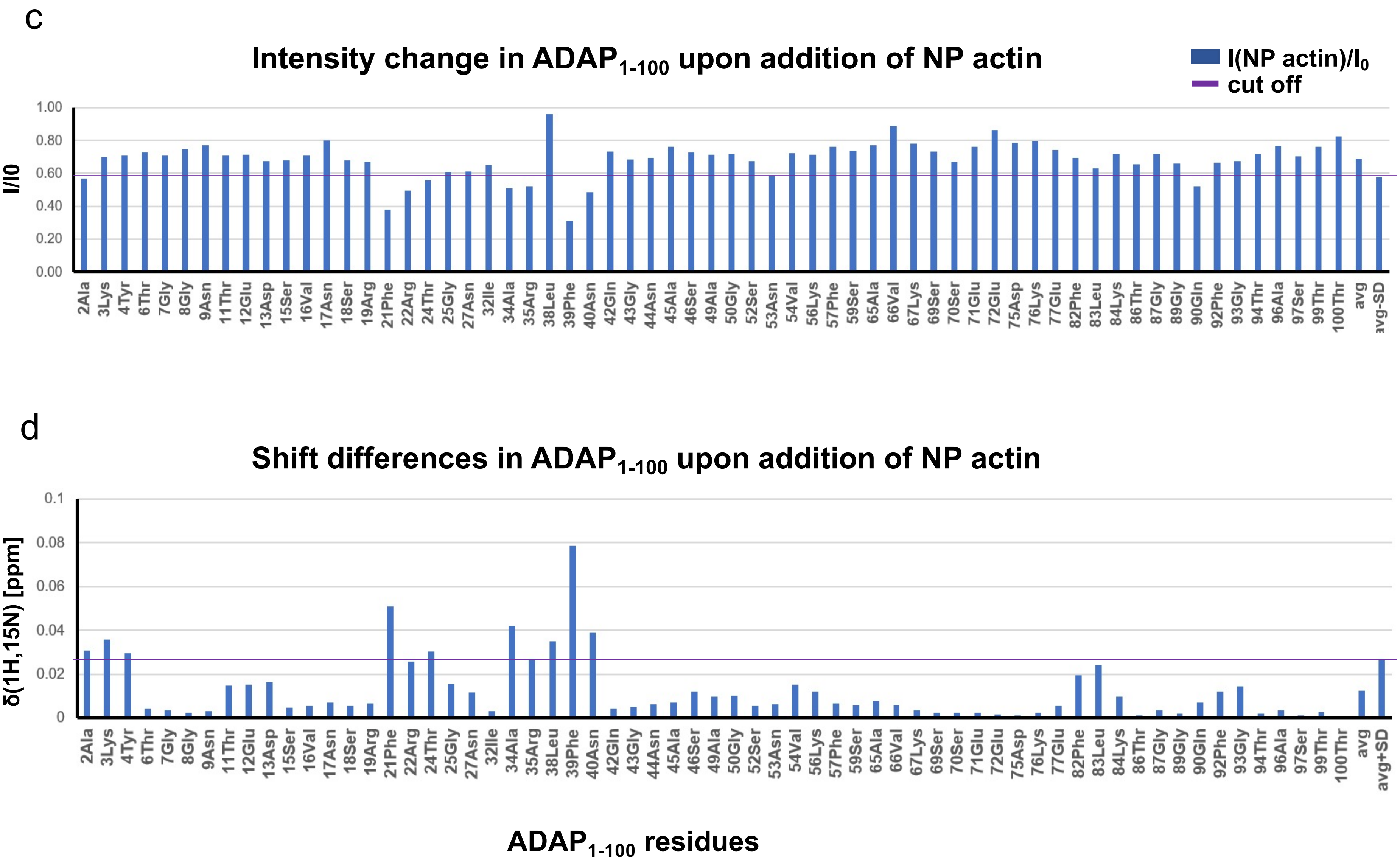

Extended Data Fig. 6

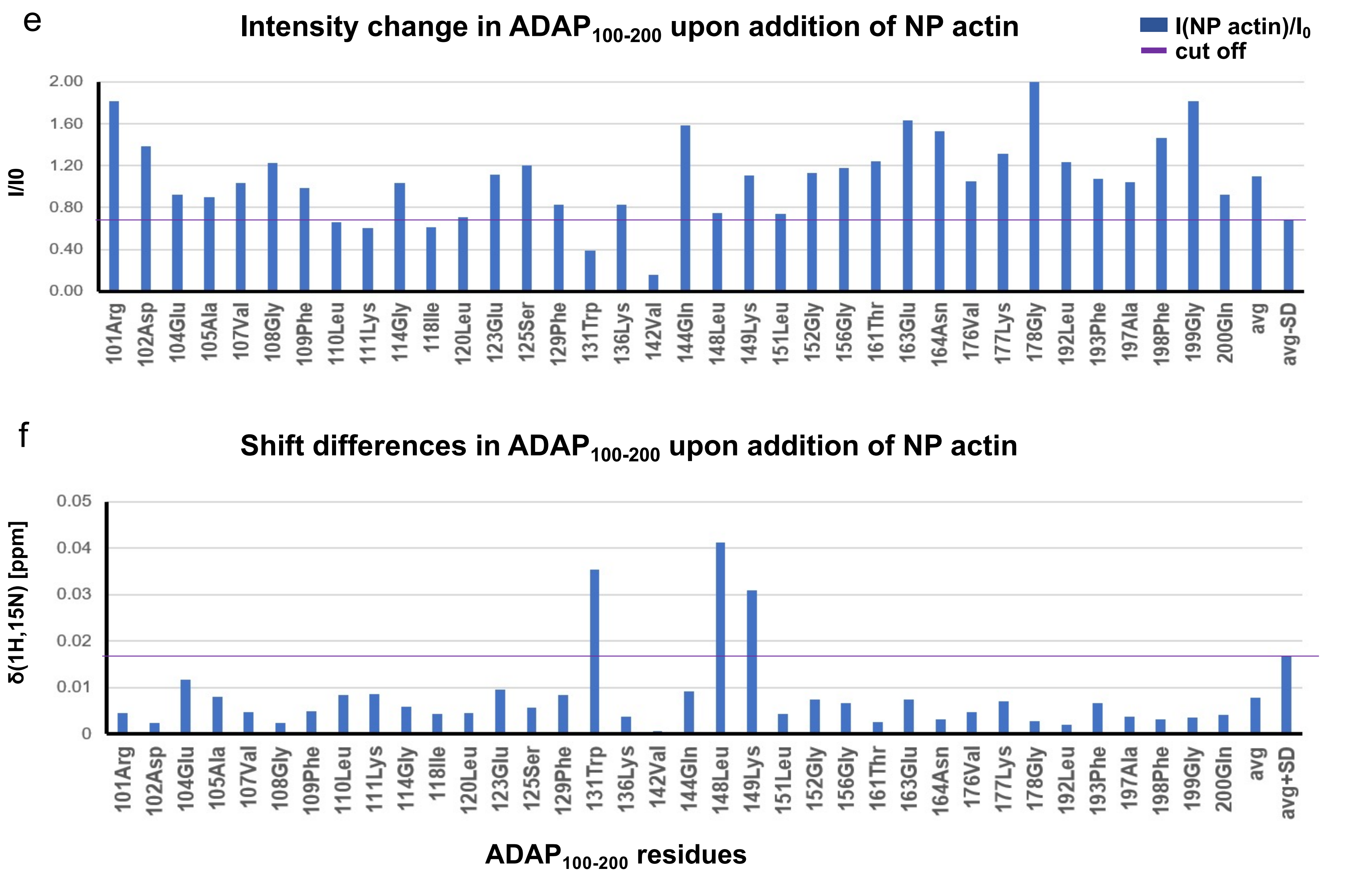

**Extended Data Fig. 6: ADAP$_{1-100}$ and ADAP$_{100-200}$ interact with monomeric actin confirmed by NMR.** (a) $^{1}H$-$^{15}N$-HSQC spectrum of deuterated, $^{15}N$-labeled ADAP$_{1-381}$ in 10mM HEPES, 50mM NaCl, 0.4mM ATP, pH 6.5 shows narrow signal dispersion which indicate disordered nature of protein. (b) Theoretical prediction of structure of ADAP$_{1-381}$ using (https://st-protein.chem.au.dk/odinpred)[47], suggests that the N-terminus of ADAP is predicted to be mostly disordered (regions highlighted in red) except for two regions (highlighted in blue). (c) The graph shows the change in $^{1}H$-$^{15}N$ peak intensity in ADAP$_{1-100}$ (I0) upon addition of NP actin (I). (d) Chemical shift changes upon addition of NP actin plotted for all assigned residues in ADAP$_{1-100}$. Both the graphs indicate major changes in the regions of residues 21-40. (e) The graph shows the change in intensity for all assigned residues in the $^{1}H$-$^{15}N$-HSQC spectrum of ADAP$_{100-200}$ (I0) upon addition of NP actin (I). (f) Chemical shift changes upon addition of NP actin plotted for all assigned residues in the $^{1}H$-$^{15}N$-HSQC spectra of ADAP$_{100-200}$. The NMR intensity change analyses indicate major changes are observed for motifs 110-111, 131, 148.149. (The analyses of spectra were performed with ccpNMR Analysis v2.4)[41].

Extended Data Fig. 7

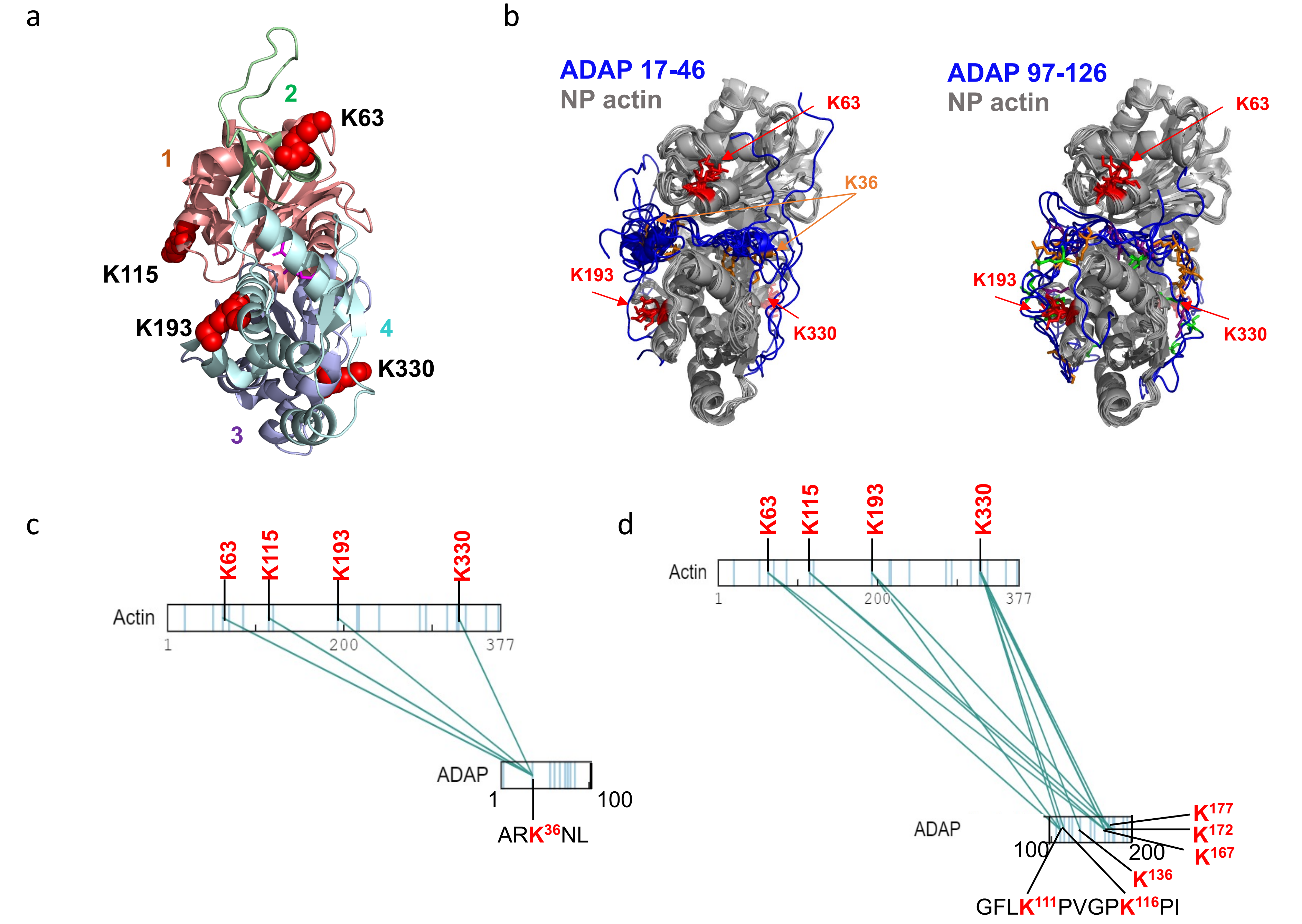

Extended Data Fig. 7

e

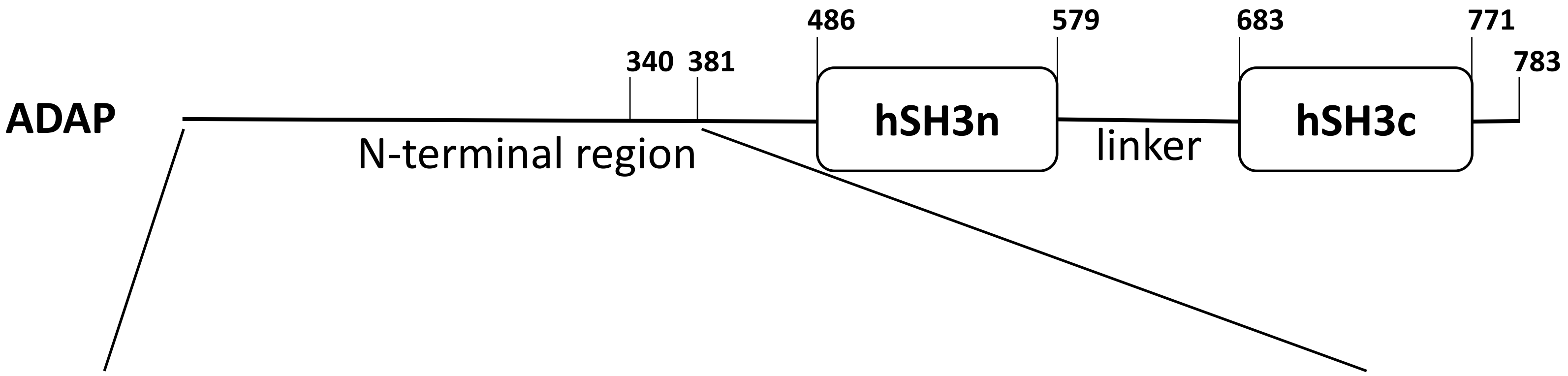


1___MAKYNTGGNPTEDVSVNSRPFRVTGPNSSSGIQARKNLFNNQGNASPPAG_50
51__PSNVPKFGSPKPPVAVKPSSEEKPDKEPKPPFLKPTGAGQRFGTPASLTT_100
101_RDPEAKVGFLKPVGPKPINLPKEDSKPTFPWPPGNKPSLHSVNQDHDLKP_150
151_LGPKSGPTPPTSENEQKQAFPKLTGVKGKFMSASQDLEPKPLFPKPAFGQ_200
201_KPPLSTENSHEDESPMKNVSSSKGSPAPLGVRSKSGPLKPAREDSENKDH_250
251_AGEISSLPFPGVVLKPAASRGGPGLSKNGEEKKEDRKIDAAKNTFQSKIN_300
301_QEELASGTPPARFPKAPSKLTVGGPWGQSQEKEKGDKNSATPKQKPLPPL_350
351_FTLGPPPPKPNRPPNVDLTKFHKTSSGNSTS_381

**(L,P)KP-motifs** **KP-motifs** **K** **R** **SKAP55 binding site**

**Extended Data Fig. 7: Crosslinking MS confirms the interaction between ADAP and monomeric actin.** (a) Crosslinked lysine residues are indicated as red spheres on subdomains of actin monomer (pdb:3hbt[45] (color code, domain 1: salmon, domain 2: green, domain 3: light-blue, domain 4: pale-cyan, ATP: magenta)**.**
(b) Docking model of ADAP peptides, which cover regions affected in NMR and cross-linking MS to G-actin. The 10 best docking models of ADAP residues 17-46 and of 97-126 are shown in blue with the structure of actin in grey (PDB 3hbt)[45]. The CABS-dock server[42] was used for docking applying the following soft 20Å distance restraints based on crosslinking MS results obtained with $ADAP_{1\text{-}381}$: ADAP K36 (orange) to each of actin residues K63, K193 and K330 (red); ADAP K106 (green) and actin K193 and K330 (red); ADAP K111 (orange) and actin K193 (red); ADAP K116 (purple) and actin K63, K193, K330 (red).
(c,d) Cross-links between $ADAP_{1\text{-}100}$ (c) or $ADAP_{100\text{-}200}$ (d) and NP-actin are shown as green lines while the cross-linked lysine residues are marked in red.
(e) Composition of the N-terminal region of ADAP highlighting K and R residues as well as KP and (L,P)KP motifs within the sequence. The SKAP55 binding site is underlined.

Extended Data Fig. 8

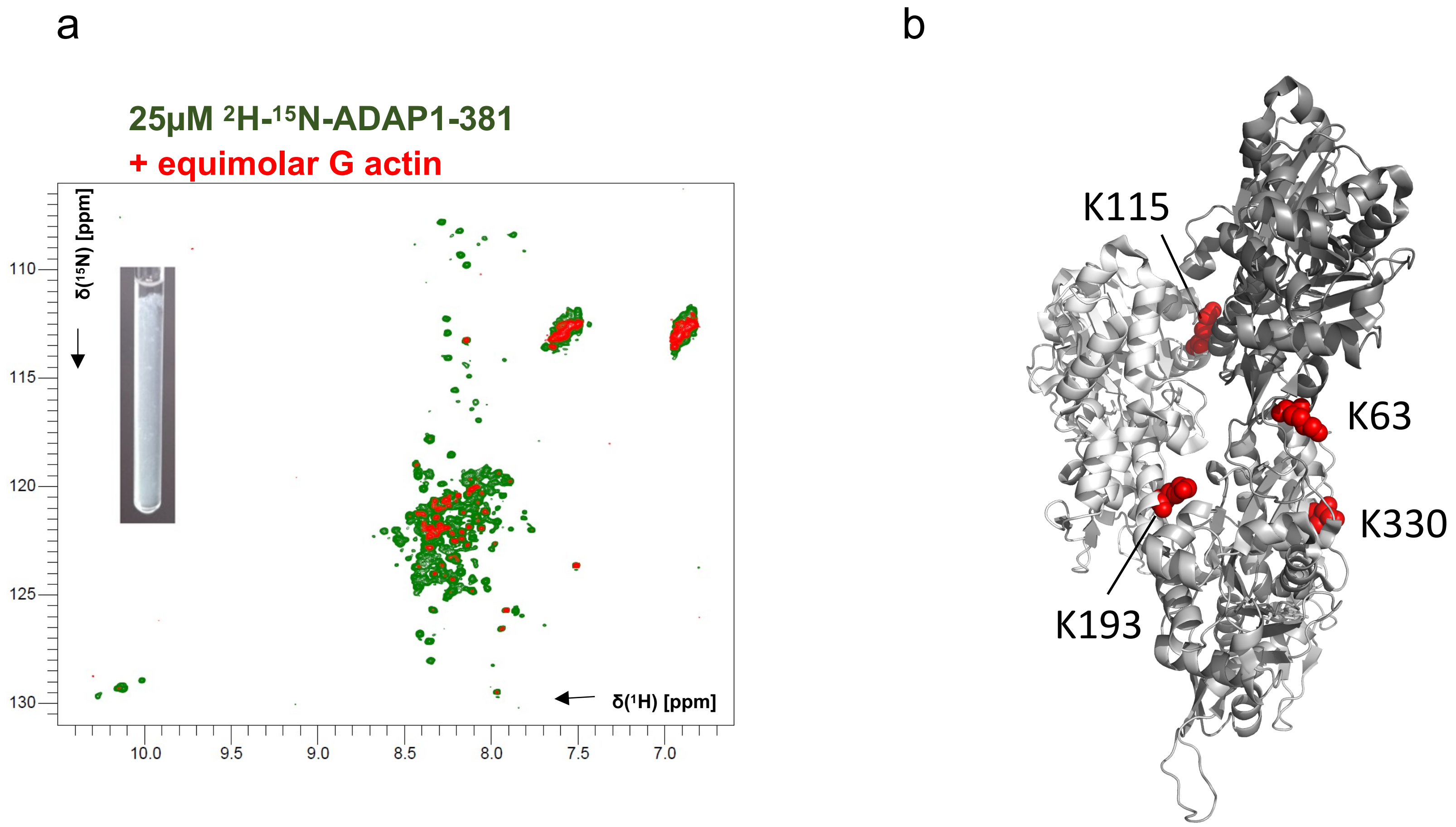

### Extended Data Fig. 8

c

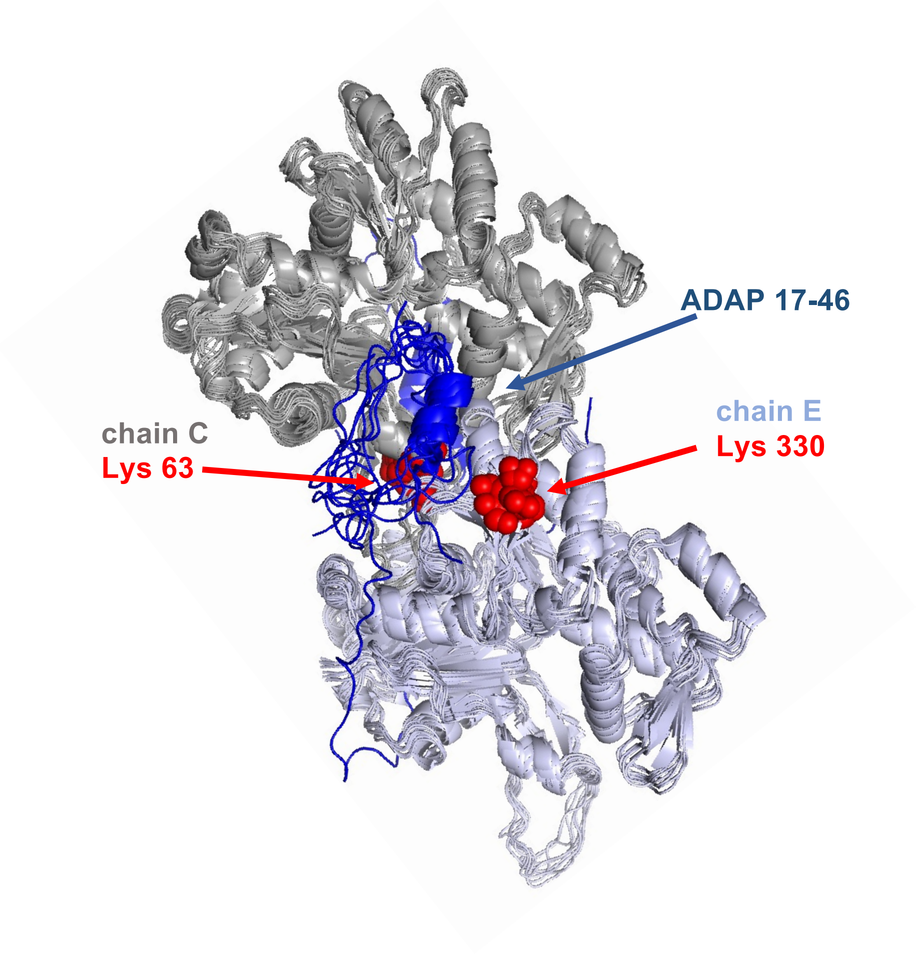


d

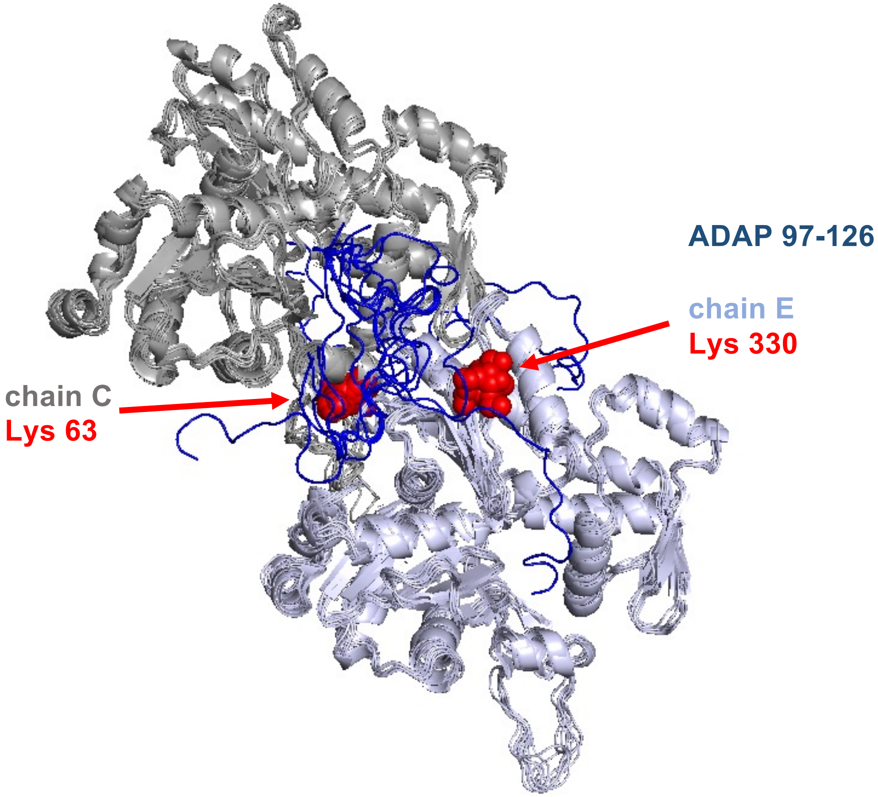

Extended Data Fig. 8

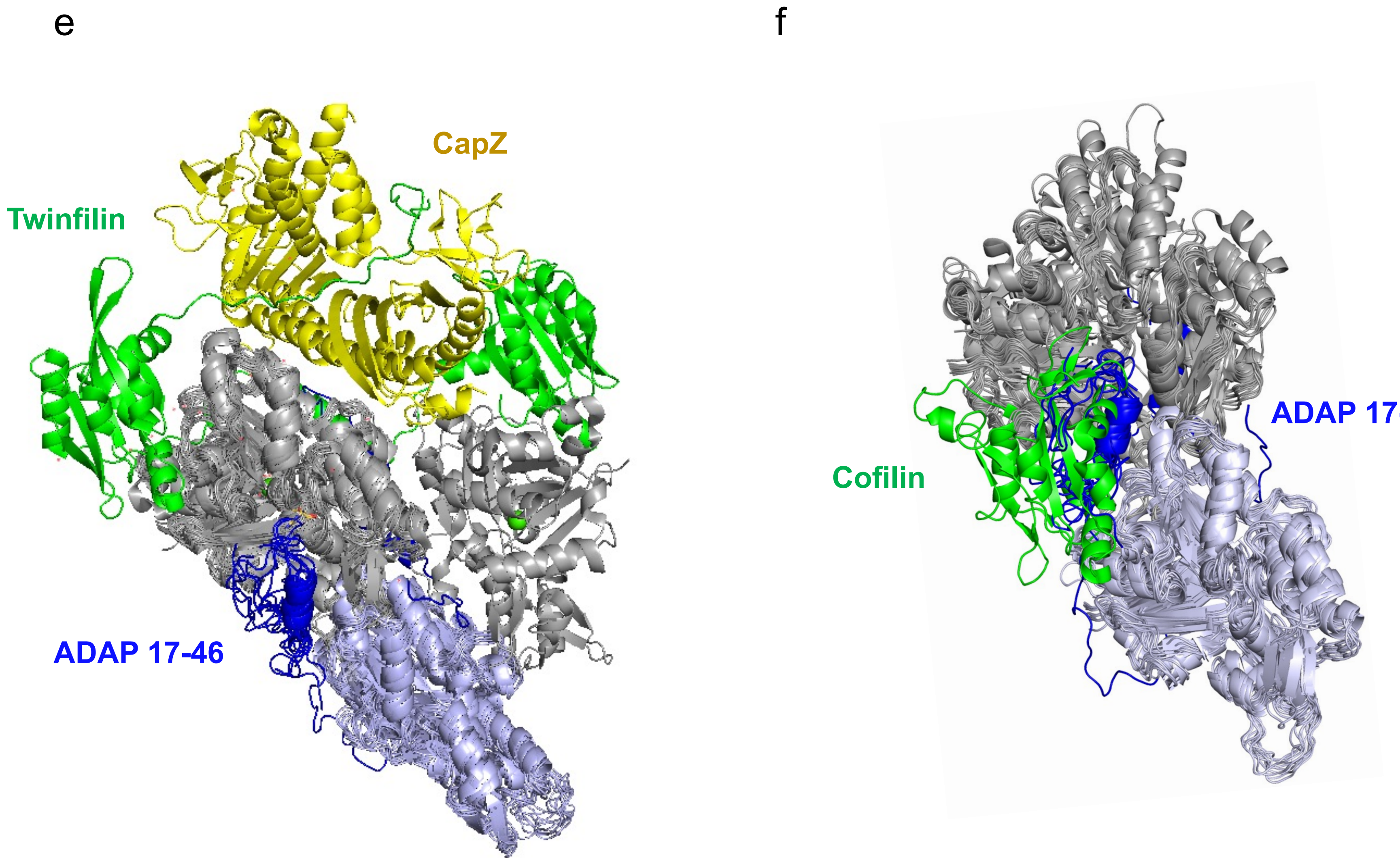

**Extended Data Fig. 8: Interaction between ADAP and filamentous actin confirmed by NMR and crosslinking MS.** (a) $^{1}H$-$^{15}N$ HSQC spectra of $ADAP_{1-381}$ (green) mixed with recombinantly expressed G-actin (red). The inset showing G-actin immediately turned turbid when mixed with $ADAP_{1-381}$ in the NMR tube. The line broadening and vanishing of peaks in the spectrum indicate strong binding with F-actin. (b) Lysine residues that cross-link to pre-polymerized F-actin are indicated as red spheres on actin dimer chosen from F-actin (chain C and chain E from pdb:6fhl[45]) (color code, domain 1: salmon, domain 2: green, domain 3: light-blue, domain 4: pale-cyan, ATP: magenta). c, d) The 10 best docking models of ADAP residues 17-46 (c) and 97-126 (d) are shown in blue with the structure of actin dimer in grey and light blue (chain C and E of PDB 6fhl). The CABS-dock server was used for docking using a 20Å soft distance restraint between K36 or K116 of ADAP and K63 of chain C and K330 of chain E of actin. (e) Structure of twinfilin (shown in green)/CapZ (shown in yellow) (PDB:7ccc)[25] bound to actin superimposed with the one model of the ADAP 17-46 bound actin dimer. (f) Structure of cofilin (shown in green, PDB: 6vao)[26] bound to actin dimer superimposed with the ten best models of ADAP residues 17-46 bound to the actin dimer showing that the binding sites overlap.
