## Supplementary_Info_Movie_References for "ADAP’s intrinsically disordered region is an actin sponge regulating T cell motility"

**Supplementary Movie 1**: Motility of ADAP wt and KO cells on pMBMECs. Movie starts after onset of physiological flow strength (1.5 dyn/cm2). Images have been acquired with a 20x objective (LD Plan-Neofluar 20x/0.4 from Zeiss) at an AxioObserver.Z1 microscope (Zeiss) every 20 seconds and the movie has been assembled using the 3 channels merged into one image each with 6 frames per second (1 second of movie corresponds to 2 minutes real time). Blue, wildtype T cells (CMAC labelled). Red, ADAP knockout T cells (CMTMR labelled). Greytones, monolayer of endothelial cells (unlabeled). Scale bar, 50 micrometer. More ADAP ko T cells (red) than wt T cells (blue) detach over the period of observation from the endothelial surface.
